# Inhibition of Retinoic Acid Signaling in Proximal Tubular Epithelial cells Protects against Acute Kidney Injury by Enhancing Kim-1-dependent Efferocytosis

**DOI:** 10.1101/2023.06.15.545113

**Authors:** M. Yang, L.N. Lopez, M. Brewer, R. Delgado, A. Menshikh, K. Clouthier, Y. Zhu, T. Vanichapol, H. Yang, R. Harris, L. Gewin, C. Brooks, A. Davidson, M.P. de Caestecker

## Abstract

Retinoic acid receptor (RAR) signaling is essential for mammalian kidney development, but in the adult kidney is restricted to occasional collecting duct epithelial cells. We now show there is widespread reactivation of RAR signaling in proximal tubular epithelial cells (PTECs) in human sepsis-associated acute kidney injury (AKI), and in mouse models of AKI. Genetic inhibition of RAR signaling in PTECs protects against experimental AKI but is associated with increased expression of the PTEC injury marker, Kim-1. However, Kim-1 is also expressed by de-differentiated, proliferating PTECs, and protects against injury by increasing apoptotic cell clearance, or efferocytosis. We show that the protective effect of inhibiting PTEC RAR signaling is mediated by increased Kim-1 dependent efferocytosis, and that this is associated with de-differentiation, proliferation, and metabolic reprogramming of PTECs. These data demonstrate a novel functional role that reactivation of RAR signaling plays in regulating PTEC differentiation and function in human and experimental AKI.

**Graphical abstract:** 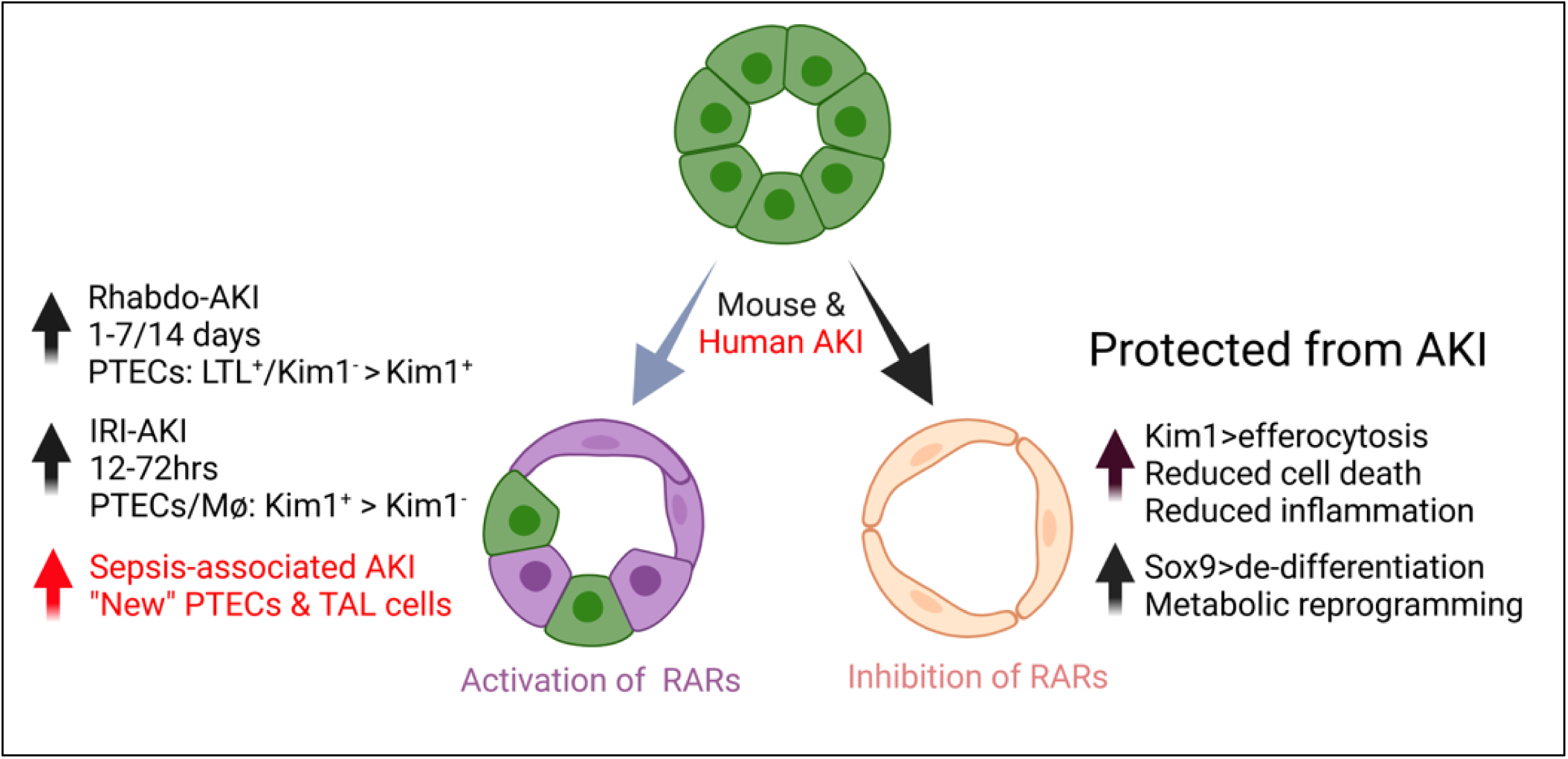

## Introduction

Acute Kidney Injury (AKI), which occurs in 10-20% of adults hospitalized in developed countries with acute illnesses, (1) is an independent predictor of mortality (2), and increases the risk of chronic kidney disease (CDK) and end stage renal disease (ESRD)(3, 4). Current treatment options for AKI are limited, and no therapeutic interventions have been shown to improve outcomes after AKI (5, 6). In part this is because AKI is a heterogeneous condition that is defined clinically based on rapid changes in renal function that does not reflect the diversity pathophysiological responses to the injury(7). The most common form of AKI is acute tubular injury (ATI), which can be caused by nephrotoxic drugs, reduced renal perfusion leading to renal ischemia, muscle injury leading to rhabdomyolysis-AKI (rhabdo-AKI), trauma and surgery including cardiac surgery, and sepsis-associated AKI (SA-AKI)(7, 8). Molecular analyses of human kidneys from patients with AKI have begun to elucidate pathophysiological mechanisms (9-11), but limited access to tissues will continue to hamper our ability to identify molecular mechanisms of AKI (12). An experimental model that has been used extensively to evaluate the cellular responses over time is the rodent ischemia reperfusion induced model of AKI (IRI-AKI)(13). This induces defined injury with dominant damage to proximal tubular epithelial cells (PTECs), the most abundant and metabolically active cells in the kidney(14). Other models of AKI used to evaluate AKI mechanisms, include cisplatin-AKI (CP-AKI), modeling the effects of chemotherapy in patients with solid organ malignancies(15), and rhabdo-AKI, modeling the effects of crush injuries(16), and heme-induced AKI in sepsis, sickle cell disease, and cardiac surgery (17, 18).

KIM-1 (encoded by *HAVCR1*) is a phosphatidylserine (PS) receptor first described as an marker of injury in de differentiated, proliferating PTECs after IRI-AKI (19). Kim-1 has since been shown to be rapidly upregulated in PTECs in a variety of rodent models and patients with AKI (20-23). Increased urinary Kim-1 is also a biomarker of tubular injury in patients with different renal diseases (21, 24, 25). *In vitro* studies indicate that Kim-1 triggers efferocytosis of apoptotic cells by PTECs (26, 27), while loss of function studies using *Havcr1* mutant mice have shown that loss of Kim-1 expression or PS binding functions, exacerbates ATI, increase tubular cell apoptosis and necrosis, and increase damaging inflammatory responses after IRI- and CP-AKI (28-30). Decreased inflammation is mediated by decreasing pro inflammatory secondary necrosis of apoptotic cells that have not undergone clearance (31), and by the suppression of inflammatory responses by PTECs that have phagocytosed apoptotic cells (29). These findings are supported by the observation that over-expression of a synthetic efferocytosis receptor in PTECs reduces inflammation and protects against IRI-AKI (32). However, in other settings, over-expression of Kim-1 may provoke inflammation and fibrosis. For example, a conditional mouse mutant that induces persistent Kim-1 expression in renal epithelium starting during embryonic kidney development, provokes renal inflammation and fibrosis in adult mice (33). In addition, persistent Kim-1 expression is a feature of pro-inflammatory PTECs that have failed to undergo productive repair after AKI (10, 34, 35). On this basis, it is unclear whether enhancing Kim-1 expression after AKI would be protective, or would cause more severe injury and CKD.

The retinoic acid (RA) signaling pathway plays an essential role in mammalian kidney development (36), but in the adult kidney, RAR signaling is restricted to occasional cells within the collecting duct (CD) system (37). RA metabolites are generated in a two-step process that includes the rate limiting and irreversible oxidation of retinaldehyde to RA by retinaldehyde dehydrogenases (RALDHs, encoded by the *ALDH1a* gene family). Once generated, RAs may act in an autocrine or paracrine fashion, and mediate their effects primarily by binding to the RA receptors (RARs), which activate or repress transcriptional targets to regulate a variety of cellular functions. These effects are cell type and context dependent. For example, RA can promote epithelial differentiation (e.g.: pancreatic and ovarian cancer cells, non malignant mammary epithelial cells, and glomerular podocytes) and enhance stem cell-like properties in other cell types (e.g.: glioma stem cells, human induced pluripotent stem cells, breast cancer cells) (38-40). In other contexts, activation of RAR signaling promotes heart, limb, and fin appendage regeneration in amphibia and fish (41-43).

We have previously shown that Raldh2 and 3 are induced in peritubular monocyte/macrophages and myofibroblasts after IRI-AKI, and have used RAR reporter mice to show that signaling is activated in adjacent renal macrophages and in injured, Kim-1^+^ PTECs after IRI-AKI (44). Systemic inhibition of RAR signaling worsens renal injury and increases inflammatory macrophage activation after IRI-AKI, and macrophage-depletion studies showed that worsening injury after inhibition of RAR signaling is dependent on macrophages (44). These findings are consistent with the known anti-inflammatory effects of RAR signaling in macrophages (45-49), and the protective effects of exogenous retinoids on renal injury in toxin and IRI-AKI models (44, 50, 51), in which RA treatments reduce both tubular injury and inflammation in the kidney (44, 50). However, these findings do not explain the functional role that activation of RAR signaling in PTECs plays in the cellular responses to injury, nor whether this response to injury is conserved in patients or in other experimental models of AKI.

In these studies, we show for the first time that PTEC RAR signaling is activated in patients with severe SA-AKI, and in mice with rhabdo-AKI, and unexpectedly show that selective inhibition of RAR signaling in PTECs protects against both IRI- and rhabdo-AKI by increasing Kim-1-dependent efferocytosis by PTECs. Increased Kim-1 expression and phagocytic activity is not associated with increased tubular injury, but is associated with enhanced de-differentiation and metabolic reprogramming of PTECs. These findings provide the first gain-of-function evidence that enhanced Kim-1 expression by injured PTECs has a protective role in AKI, and suggest that activation of RAR signaling after AKI is a compensatory response that acts to promote or maintain the mature differentiation state and quiescence of surviving PTECs after AKI.

## Results

### Retinoic acid signaling is activated in proximal tubular epithelial cells in human and experimental AKI

To determine whether RAR signaling is activated in human AKI, we evaluated bulk RNA sequencing (RNA Seq) and single nuclear RNA seq (snRNA Seq) data obtained from a validated cohort of samples from patients with severe SA-AKI (see S. Methods for a summary of the clinical and validation criteria used for these studies) (52). Gene set enrichment analysis (GSEA) was performed to assess enrichment with 27 validated RAR target genes, (26 upregulated, and 1 variably regulated by RA in different cell types) (53), and 3 RA regulatory genes, *Aldh1a2*, *Aldh1a3*, and *Cyp26B1*, that are upregulated in mouse kidneys after IRI-AKI (S. Table 1) (44). There was enrichment for RAR target genes in bulk and PTEC RNA seq datasets (Fig. 1A-C, S. Table 2), while thick ascending limb (TAL) and renal leukocyte populations showed less robust enrichment for RAR target genes (Fig. 1A, S. Table 2). There was no significant enrichment for RAR target genes in collecting duct (CD), or other snRNA Seq cell populations in this dataset (Fig. 1A, S. Table 2). More detailed analysis of PTEC subsets indicates that RAR target genes are predominantly activated in injured, repairing, and failed repair PTECs (which also express Kim-1), but also in differentiated S3 segment PTECs in AKI samples (S. Fig. 1). RAR target genes are also expressed in “new” injured and repairing TAL cells (S. Fig 2), and in monocyte/macrophage lineages (S. Fig. 3).

**Figure 1.**
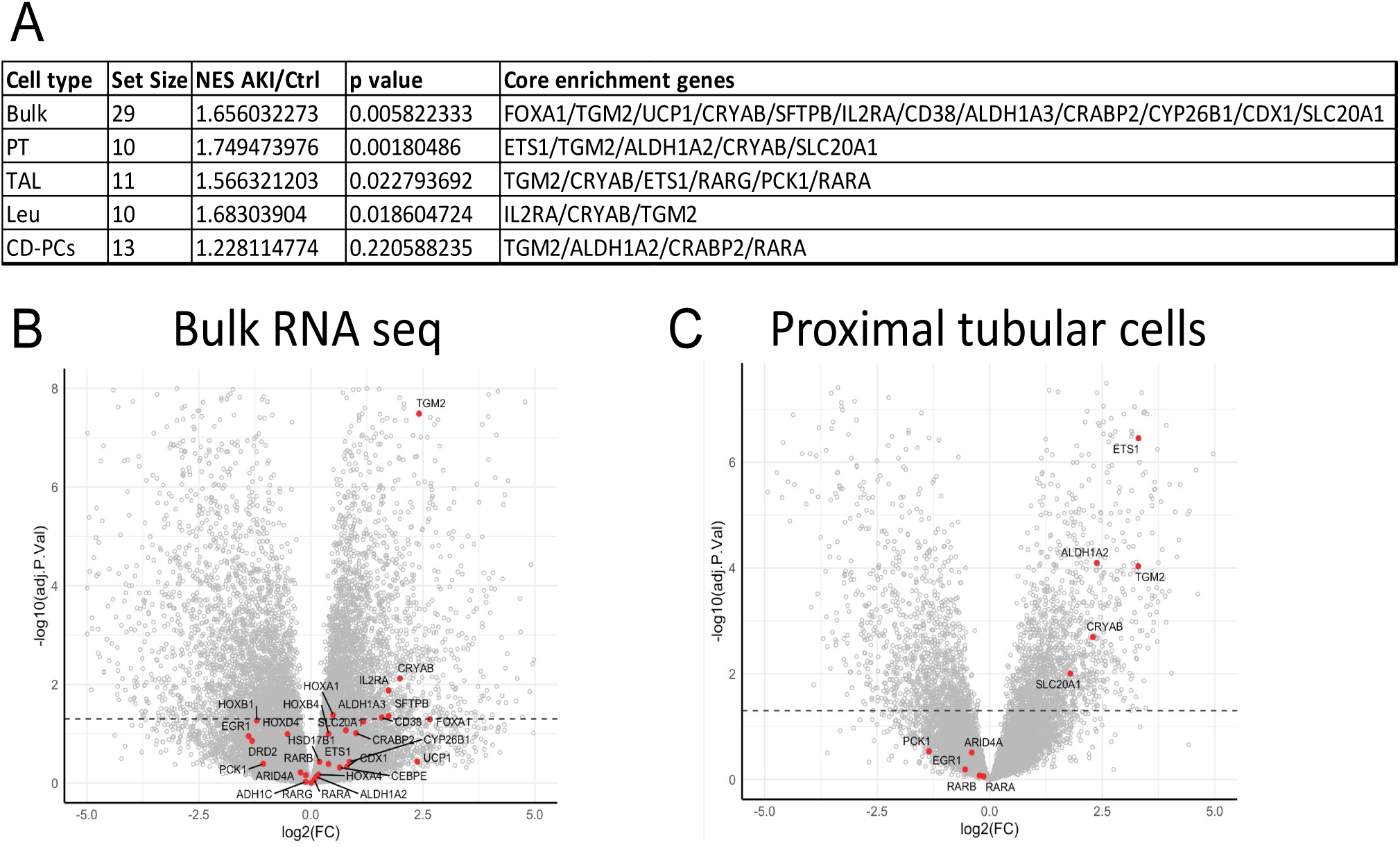
Activation of Retinoic Acid Receptor (RAR) signaling in patients with Sepsis-Associated AKI. Gene set enrichment analysis (GSEA) of bulk RNA, and cell-specific snRNA sequencing kidney datasets from patients with Sepsis-Associated AKI (SA-AKI) compared with age matched controls from Hinze et al. 2022.^1^ ***A, Increased expression of RAR target genes in kidneys from SA-AKI patients.*** GSEA for RA targets genes in the RNA seq datasets. Cell types: PT, proximal tubular epithelial cells; TAL, thick ascending limb; Leu, leukocytes; and CD, collecting duct. Set size=no. of genes from the RAR gene set that are represented in each dataset. NES=normalized expression score for the gene set size in AKI vs. control samples. p values are uncorrected since each RNA seq dataset was only compared with a one RAR target gene set; Core enrichment genes=genes within the gene set that are driving the NES. **B/C,** Volcano plots indicating fold change in expression of core enrichment gene (AKI vs. controls) from bulk RNA and PTEC snRNA seq datasets. Dotted line indicates p<0.05.

These data are consistent with our prior findings using the validated *RARE-hsp68-LacZ* (RARE-LacZ) transgenic reporter mouse line, which expresses bacterial LacZ/β-Galactosidase in cells upon activation of canonical RAR signaling (36, 37, 54). Using this reporter line, we found there was strong activation of the reporter in Kim-1^+^ PTECs and F480^+^ renal macrophages, with less activation of the reporter in THP-1^+^ TAL and Dolichos Bifluoris (DB)^+^ CD cells, after IRI-AKI (44). To determine whether there were differences in the cellular distribution of RAR activation in different models of AKI, we evaluated RAR activation in a mouse model of rhabdo-AKI induced by intramuscular injection of glycerol (55). In this model, there is upregulation of PTEC, and distal tubular injury markers, *Kim-1* and *N-Gal/Lcn2* mRNAs, respectively, and upregulation of the ubiquitous, early injury response gene, *Heme Oxygenase 1 (Ho-1)*, 24-72 hrs after AKI (Fig. 2A/B). This was associated with upregulation of the RAR target gene, *Rarb*, as well as other RAR targets and regulators that were upregulated in the bulk RNA Seq and PTEC snRNA Seq human SA-AKI datasets described above (Fig. 2C). To evaluate the cellular distribution and kinetics of RAR signaling in more detail, we evaluated the distribution of the RARE-LacZ reporter after rhabdo-AKI. We observed widespread activation of the reporter gene throughout the cortex and outer stripe of the outer medulla (OSOM) of the kidney, peaking 3 to 7 days after induction of injury and returning to baseline expression by day 14 (Fig. 2D/E). This contrasts with IRI-AKI in which the RARE-LacZ reporter activation is largely restricted to the outer medulla, and was more transient, peaking 12-24 hrs after injury, and returning to baseline levels by day 7 (44).

**Figure 2.**
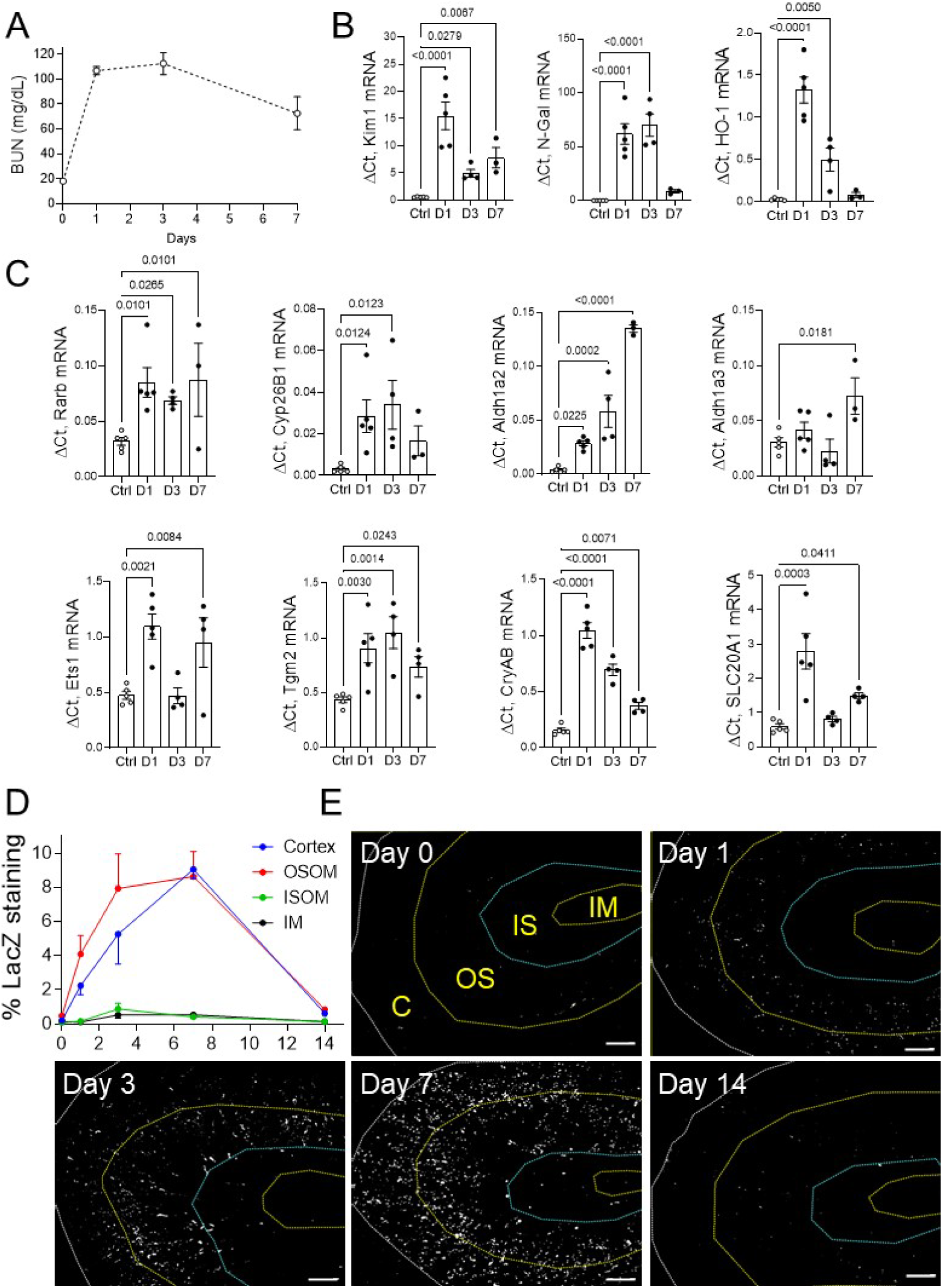
Widespread activation of RAR signaling in the kidneys of mice after rhabdomyolysis-induced AKI. BALB/c mice were injected with IM glycerol. 5 mice were euthanized before injury and at Day 1, 4 at Day 3, and 3 at Day 7 after glycerol injection. **A,** BUN time course. Days after injury on the X axis. **B,** Expression of kidney injury markers. QRT-PCR for injury markers *Kim1, N-Gal* and *HO-1* mRNAs. ***C, RAR target gene expression after rhabdo-AKI.*** RAR target genes. QRT-PCR for RAR target genes *Rarb, Cyp26B1, Ets1, Tgm2, CryAB,* and *Slc20A1*, and RA regulators, *Aldh1a2/Raldh2*, and *Aldh1a3/Raldh3* mRNAs by QRT-PCR. Results are expressed as mean +/-SEM, with individual time points indicated in B and C. One way ANOVA used to determine statistical significance, and q values are shown after correcting for repeated comparisons. ***D, Widespread and prolonged activation of RAR signaling in the kidney after rhabdo-AKI.*** Spatial distribution and kinetics of RAR signaling. RARE-LacZ reporter mice were injected with 5ml/kg of 50% glycerol into both thighs after overnight water restriction. Kidneys were harvested and frozen sections stained for LacZ activity, 3 before injury (Day 0); 4 at Day 1 and 3; 3 at Day 7; and 5 at Day 14 after injury. The % of total areas staining for LacZ activity in the cortex (C); outer stripe of the outer medulla (OSOM); inner stripe of the outer medulla (ISOM); and inner medulla (IM), as indicated in E. E, Representative images showing LacZ staining at different time points after injury. LacZ staining is pseudo-colored in white, and regions of the kidney are demarcated by dotted lines in the first panel. Scale bars, 500μM.

To explore the cellular localization of RAR signaling after rhabdo-AKI, we co-stained the RARE-LacZ reporter mouse kidneys for bacterial LacZ/β-Galactosidase and tubular segment and inflammatory cell markers. The majority of LacZ^+^ cells were localized in LTL^+^ PTECs in the cortex and OSOM (Figs. 3A/B). Less than 1% of LacZ^+^ cells were Kim-1^+^ PTECs 24hrs after injury, increasing to 17% in the OSOM, and 21% in the cortex 72 hrs after injury (Fig. 3C/D). This contrasts with our published data in mice after IRI-AKI (44), where greater than 40% of LacZ^+^ cells were localized in Kim-1^+^ tubules in the OSOM 24hrs after injury (S. Fig. 4). Compared to PTECs, there was a lower proportion of LacZ^+^ cells in THP-1^+^ TAL cells in the cortex, OSOM, and the inner stripe of the outer medulla (ISOM) (S. Fig. 5A/B), and in F4/80^+^ renal macrophages in the OSOM after rhabdo-AKI (S. Fig. 5C/D). In addition, there were only a small number of LacZ^+^ cells in AQP2^+^ CD cells 1 to 7 days after rhabdo-AKI (S. Fig. 5E/F). The % of LacZ^+^ AQP2^+^ CD cells increased in the OSOM by day 14 (S. Fig. 5E/F). However, at this time point, the total numbers of LacZ^+^ cells in the kidney have returned to near baseline levels (Fig. 2D/E), so that the total number of AQP2^+^ CD cells with activation of RAR signaling 14 days after injury, is low. These findings are consistent with human data showing dominant activation of RAR signaling in PTECs from patients with SA-AKI, but also indicate there are differences in the kinetics and localization of RAR signaling in different AKI models.

**Figure 3.**
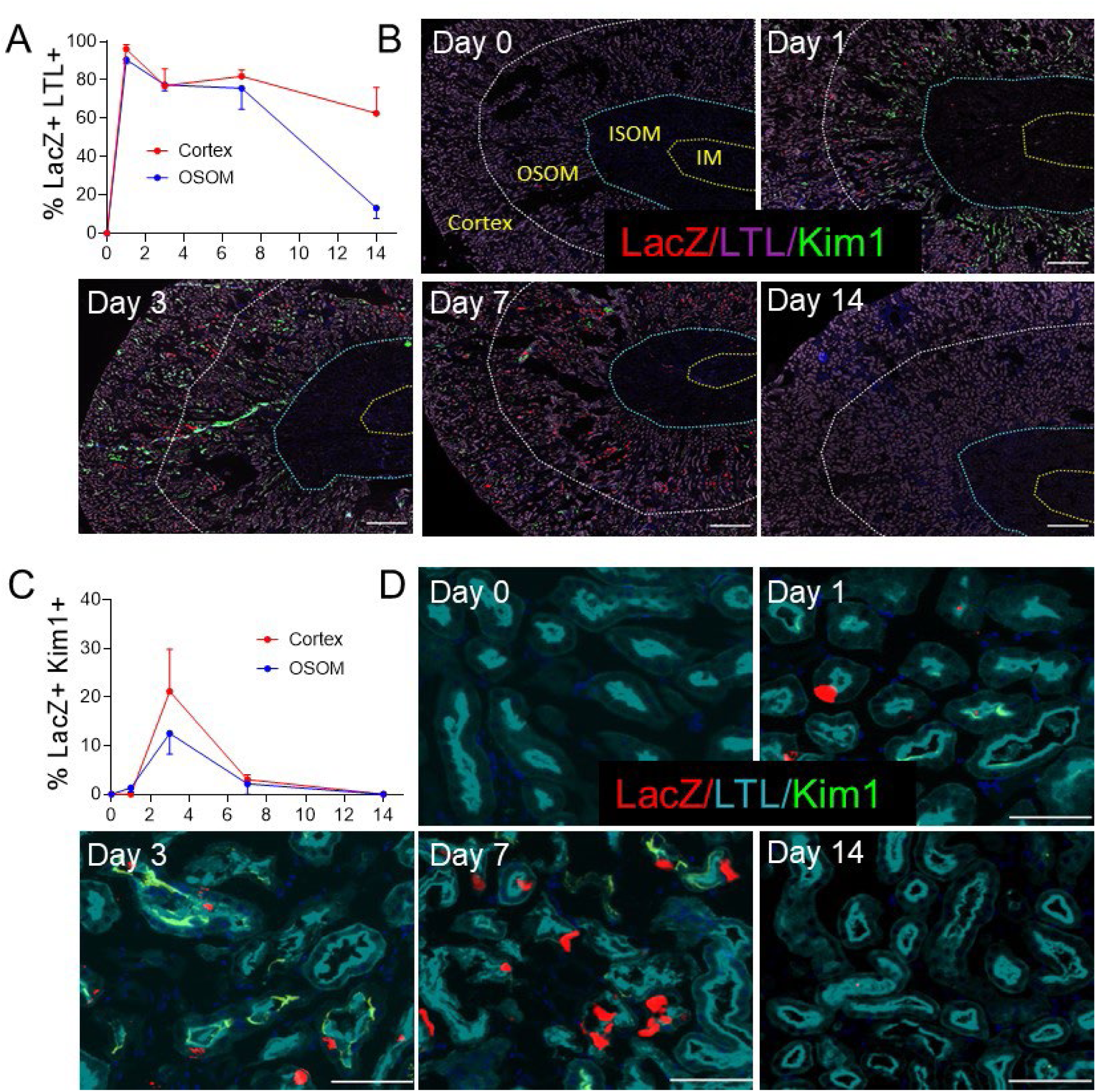
RAR signaling is extensively activated in Kim-1 negative PTECs after Rhabdomyolysis-AKI. RARE LacZ reporter mice were used to evaluate the cellular localization of RAR signaling after Rhabdo-AKI. After LacZ staining, kidney sections were stained with Lotus Tetraglonolobus lectin (LTL to identify PTECs), and antibodies to Kim1 (injured PTECs). ***A/C, RAR signaling is dominantly activated in LTL^+^ Kim-1^-^ PTECs.*** The % LacZ^+^ cells in LTL^+^ and Kim-1^-^ PTECs **(A)**, and in Kim-1^+^ PTECs **(C)** at the different time points after injury in the cortex and OSOM. Results expressed as means +/- SEM for 2 mice before injury (Day 0), and 4 each at Days 1, 3, 7 and 14 after injury. **B/D**, Representative images showing LacZ localization in PTECs after injury. LacZ, LTL and Kim-1 staining, as indicated. Low power images of the kidneys, regions of the kidney are demarcated by dotted lines, as indicated. Scale bars, 500μM (B). High power images of the OSOM. Scale bars, 100μM (D).

### Inhibition of retinoic acid signaling in proximal tubular epithelial cells protects against acute tubular injury

To explore the functional role of PTEC RAR signaling after AKI, we crossed *Rosa26-LSL-RARaT403X (R26R-DN RAR)* mice (36), which express a Cre-activated, truncated dominant negative RARa mutation (which inhibits RARa, b and g), with *PEPCK-Cre* mice (56), which induce efficient Cre-dependent recombination in PTECs (44). The *R26R-DN RAR* allele was bred to homozygosity to induce efficient inhibition of PTEC RAR signaling (PTEC DN RAR mice), as described (36). In previous studies we showed that PTEC DN RAR mice have normal kidneys, but have increased renal Kim-1 expression after IRI-AKI compared with *Cre-* controls (44). This was thought to reflect increased PTEC injury, but notably was not associated with increased tubular injury scores determined in PAS-stained sections. We now show that there is a non significant improvement in renal function, and an increase in *Kim-1* mRNA and Kim-1^+^ staining extending from the OSOM into the cortex of PTEC DN RAR mouse kidneys 3 days after IRI-AKI (Fig. 4A/B, E/F, S. Fig 6A/C). However, this was unexpectedly associated with reduced tubular injury, as determined by tubular injury scores in PAS-stained sections (Fig. 4C/D), and a reduction in *N-Gal/Lcn2 mRNA* expression, which is expressed by injured TAL, CD and distal tubule cells, but not PTECs (S. Fig 6A) (57). To explore this further, we evaluated the severity of rhabdo-AKI in PTEC DN RAR mice. Here we saw an improvement in renal function up to 3 days after induction of AKI (Fig. 4G/H). This was also associated with a reduction in tubular injury in the cortex and OSOM of PTEC DN RAR mice (Fig. 4I/J), and an increase in *Kim-1* expression and surface area of Kim-1 staining, but not in renal *N-Gal mRNA*, compared with *Cre-* controls (S. Fig. 6B/D, Fig. 4K/L).

**Figure 4.**
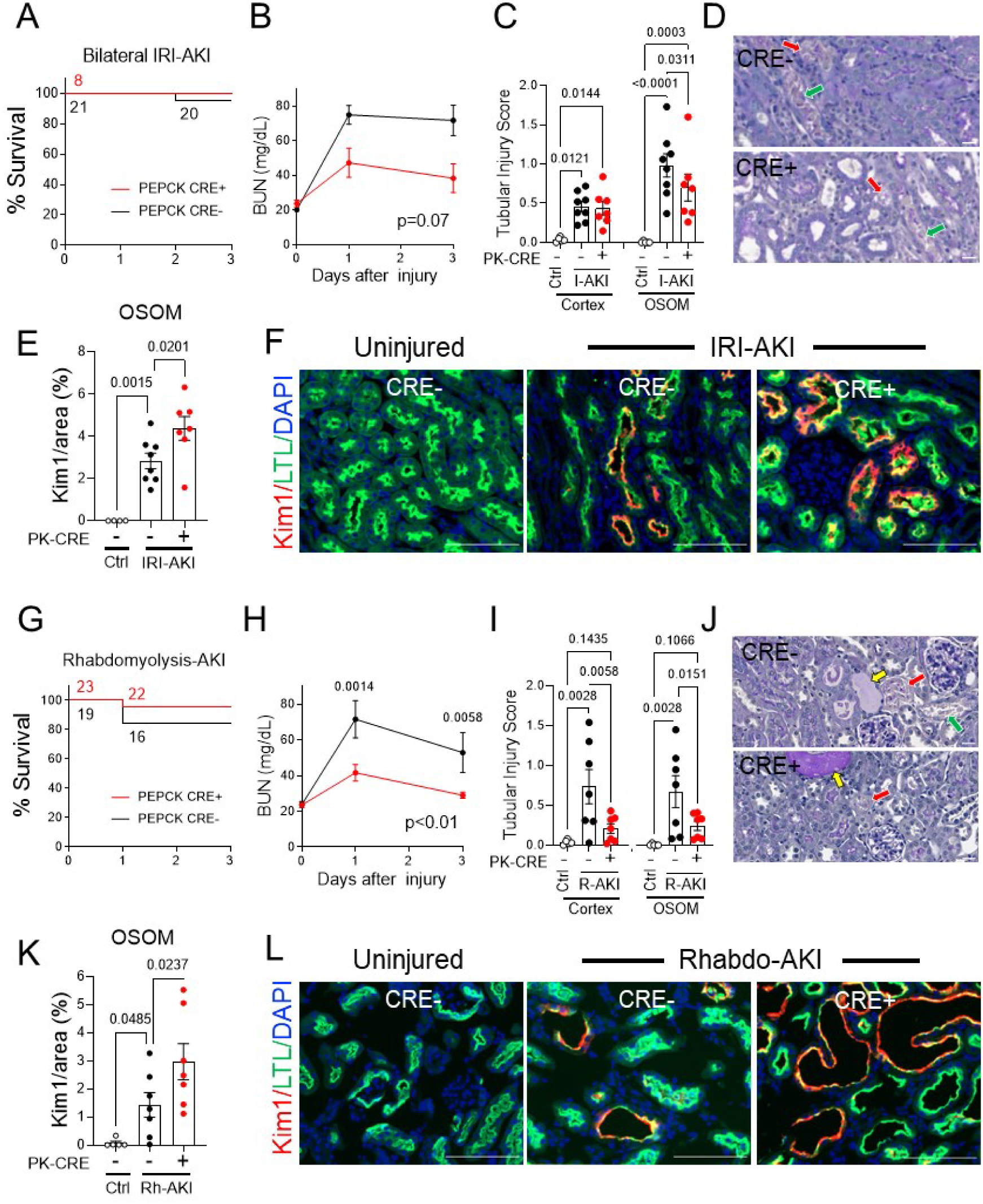
Inhibition of RAR signaling in PTECs protects against AKI but also increases Kim-1 expression in PTECs. *PEPCK Cre+ DN-RAR* (PTEC DN RAR) mice and *Cre-* controls mice underwent bilateral IRI-AKI, or rhabdo-AKI. ***A-F, Bilateral IRI-AKI.*** PTEC DN-RAR mice were monitored for 3 days after surgery. **A,** Survival curves. Mouse numbers indicated after each event. B, BUN time course after injury, mouse numbers indicated in **(A). C,** Tubular injury scores 3 days after injury in the cortex and OSOM. **D,** Representative images of PAS-stained kidney sections in the OSOM. Red arrows indicate necrosis; green arrows, detached epithelial cells. **E,** Quantification of Kim-1 staining. The % area staining with Kim-1 in the OSOM. **F,** Representative images of Kim-1 and LTL staining in OSOM. ***G-L, Rhabdomyolysis-AKI.*** PTEC DN-RAR mice were monitored for 3 days after injected with 50% glycerol. **G,** Survival curves. **H,** BUN time course, mouse numbers indicated in **(G). I,** Tubular injury scores. **J,** PAS-stained kidney sections. Red and green arrows, as described above; yellow arrows indicate tubular casts. **K,** Quantification of Kim-1 staining in the OSOM. **L,** Kim-1 and LTL staining in OSOM. Results expressed as means+/- SEM with mouse numbers indicated, or individual datapoints shown. A/G: Survival curves compared by Log-rank test, p>0.05. B/H, 2-way ANOVA to compare between groups over time. If p values <0.05 between groups, q values shown after correcting for repeated testing. C/E/I/K, 1-way ANOVA, and if p values <0.05, q values shown for between group comparisons after correcting for repeated testing. Scale bars for D/J=20μM; F/K=100μM.

To determine whether inhibition of RAR signaling in PTECs affects tissue repair after AKI, we evaluated long-term renal outcomes in PTEC DN RAR mice after IRI and rhabdo-AKI. We used a model of unilateral IRI-AKI with delayed contralateral nephrectomy (DN, Nx), which allows for the induction of severe AKI, and results in long-term fibrosis and reduced renal function (58). Using this model, we saw a significant reduction in BUN immediately after Nx in PTEC DN RAR mice, but no difference in tGFR or renal fibrosis 28 days after injury compared with *Cre-* controls (S. Fig 7A-C). Likewise, there was improved survival and reduced BUN, particularly in the first 3 days after rhabdo-AKI, but no differences in BUN or tubular injury scores 14 days after the initiating injury in PTEC DN RAR mice (S. Fig. 7D-F). These studies indicate that the principal effect of inhibiting RAR signaling in PTECs is to reduce early tubular injury, and not to reduce fibrosis or improve long-term renal function. This is also consistent with the early time course of RAR activation in PTECs, particularly after IRI-AKI.

### Inhibition of RAR signaling promotes de-differentiation and metabolic reprogramming of proximal tubular epithelial cells

Because Kim-1 expression usually correlates with the severity of tubular injury after AKI (20-23), our finding that increased Kim-1 expression was associated with reduced tubular injury in PTEC DN RAR mice was unexpected, and suggests that inhibition of RAR signaling results in the upregulation of Kim-1 expression in less injured PTECs after AKI. This is supported by the observation that there is an extension of Kim-1 staining into the cortex after IRI-AKI in PTEC DN RAR mice (S. Fig. 6C), a region of the kidney that is relatively spared from injury compared with the OSOM in this model (14). In addition to Kim-1 staining in PTEC DN RAR mice, we also found increased numbers of Sox9^+^ PTECs, a transcription factor that is expressed by de-differentiated and regenerative PTECs after AKI (59, 60), and Ki67^+^ PTECs, a marker of active cell proliferation, after rhabdo-AKI (Fig. 5A-D). In the IRI-AKI model we found a similar increase in Ki67^+^ PTECs but without an increase in Sox9 staining (Fig. 5E-G). In the rhabdo-AKI model, the majority of Sox9^+^ PTECs were also Kim-1^+^ both in *PEPCK Cre+* and *Cre-* mouse kidneys (Fig. 5B). To examine this in more detail, we evaluated the same markers in uninjured PTEC DN RAR kidneys. In the absence of injury, the % areas staining Kim-1^+^ was substantially lower than after AKI (see Fig. 4E/K). However, there was increased Kim-1 staining in the OSOM of uninjured PTEC DN RAR mice associated with increased numbers of Sox9^+^, but not Ki67^+^, PTECs compared to *Cre-* controls (Fig. 5A-D, Fig. 6A). There was also increased expression of *Kim-1* and *Sox9* mRNAs, and also increased expression of *FoxM1*, which is upregulated in de-differentiated, proliferating PTECs after IRI-AKI (Fig. 6B) (34). Scoring for histological features of ATI also suggested increased tubular injury (Fig. 6C/D). However, unlike AKI where tubular injury is characterized by the presence of tubular necrosis, detached epithelium and tubular casts (see Fig. 4D/J), in uninjured PTEC DN RAR mice, the tubular injury score was driven by the detection of de-differentiated, flattened epithelial cells, and the presence of interstitial inflammatory cells (Fig. 6D).

**Figure 5.**
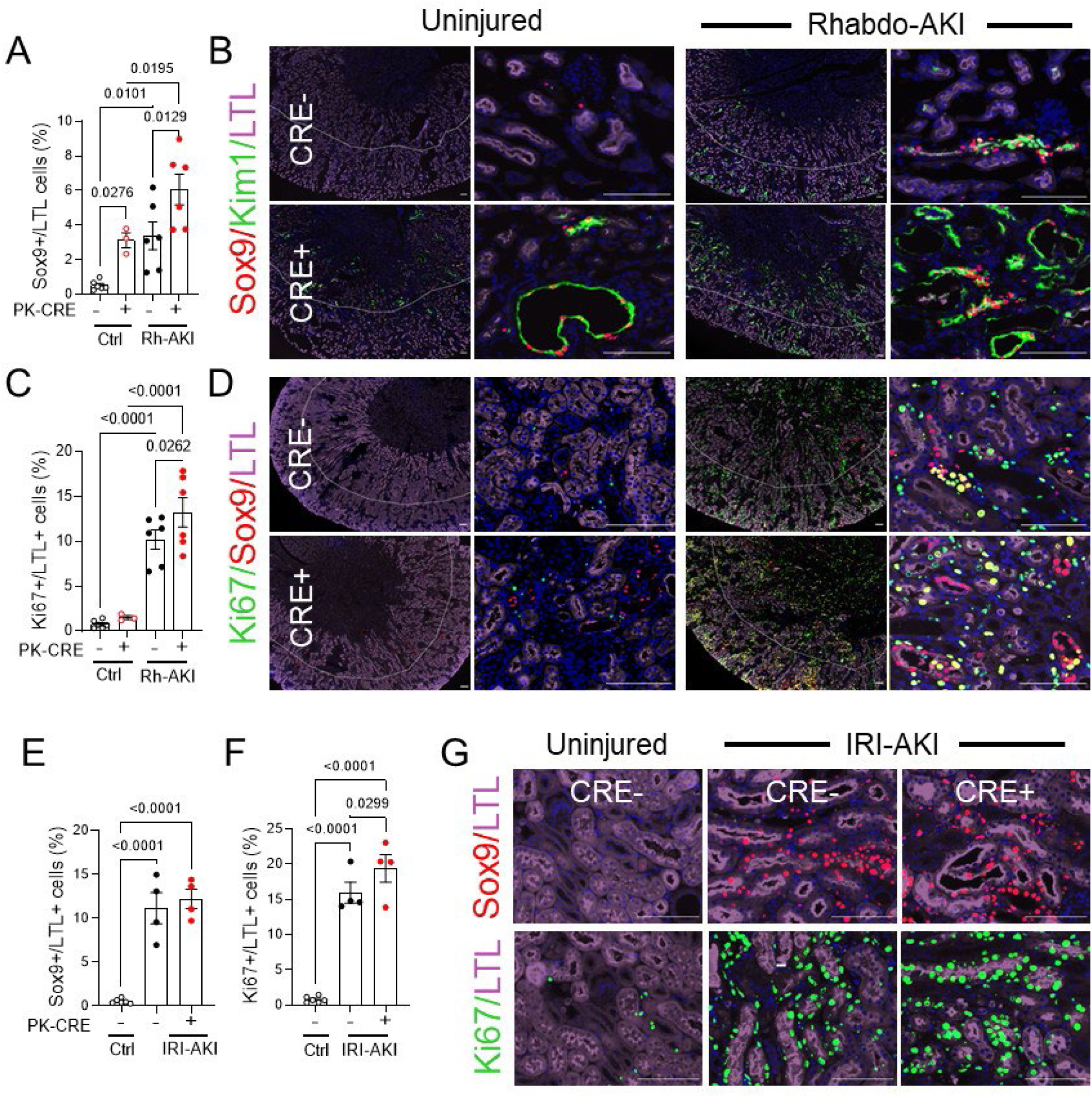
PTEC DN-RAR mice have increased expression of PTEC de-differentiation and proliferation markers after AKI. PTEC DN-RAR mice underwent rhabdomyolysis-AKI or bilateral IRI-AKI, and kidneys harvested after 3 days. ***A-D, Rhabdomyolysis-AKI***, expression of the de-differentiation marker Sox9 in PTECs. **A,** Quantification of the percentage of Sox9^+^ LTL^+^ cells in the OSOM of *PEPCK (PK)-Cre-* and *Cre+* uninjured mice (Ctrl) and 3 days after rhabdo-AKI (Rh-AKI). **B,** Representative images showing Sox9, Kim-1 and LTL staining. Left hand panels are low magnification images showing staining in the outer medulla, extending into the cortex. Right hand panels show higher magnification of the OSOM. **C,** Expression of the proliferation marker Ki67 in PTECs. Quantification of the percentage of Ki67^+^ LTL^+^ cells in the OSOM. **D,** Representative images showing Ki67, Sox9, and LTL staining. ***E-G, Bilateral IRI-AKI.* E/F,** Quantification of Sox9^+^ and Ki67^+^ LTL^+^ cells in the OSOM 3 days after IRI-AKI. **G,** Images showing Sox9 and Ki67 staining in the OSOM. Graphical results are expressed as means +/- SEM, with individual datapoints shown. 1-way ANOVA used to compare between groups, and if p values <0.05, q values shown for between group comparisons after correcting for repeated testing. Scale bars=100μM.

**Figure 6.**
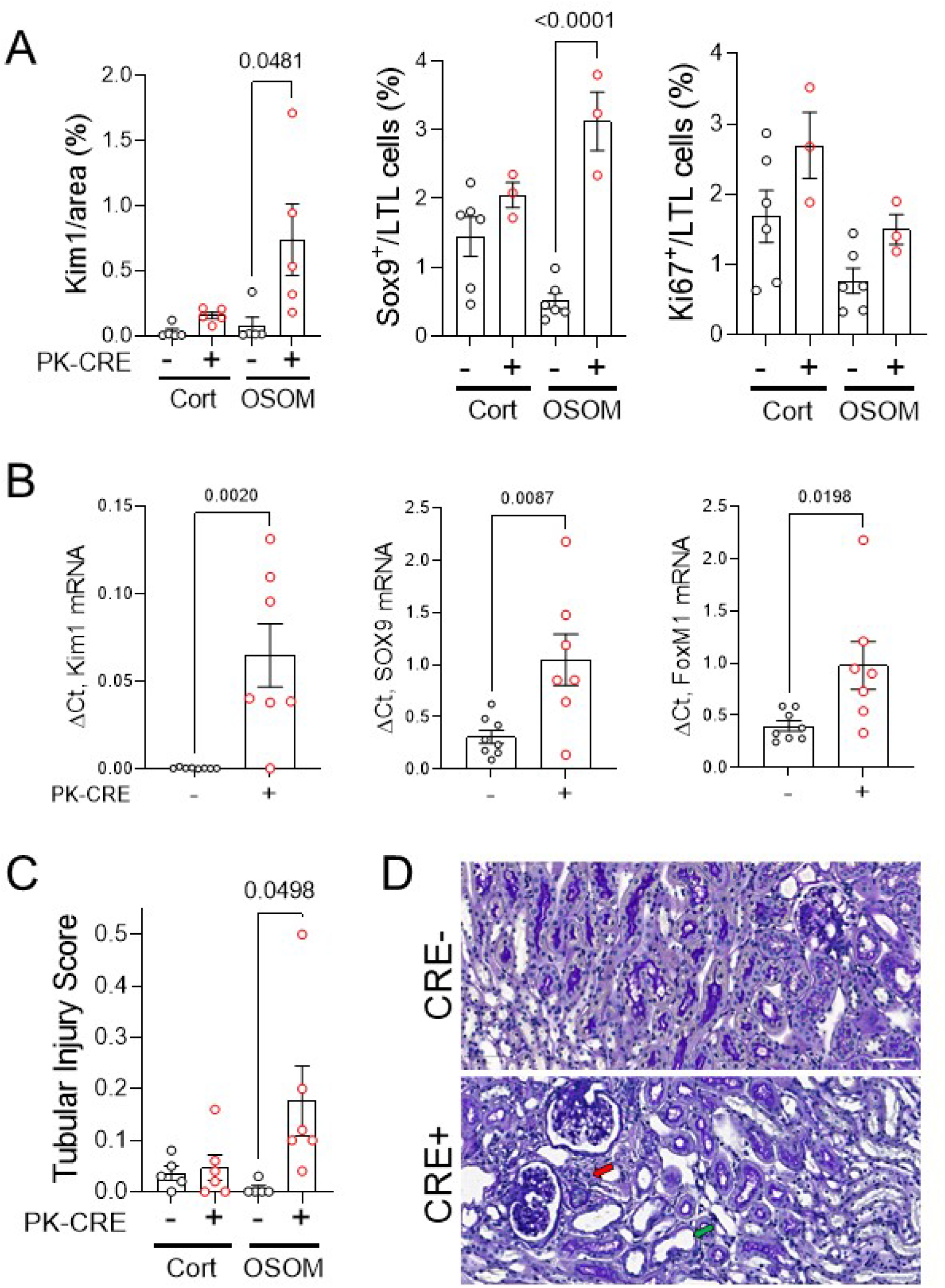
Increased expression of Kim-1 and de-differentiation markers in uninjured PTEC DN-RAR mouse kidneys. **A,** Quantification of Kim-1 (percent area stained), and the percentage of Sox9^+^ and Ki67^+^ LTL^+^ PTECs in cortex and OSOM of uninjured *PEPCK Cre+* and *Cre-* mice. Images showing Ki67, Sox9 and Kim-1 staining in uninjured *PEPCK Cre+* and *Cre-* mouse kidneys shown in Fig. 5B/D. **B,** Quantification of renal *Kim-1*, *Sox9*, and *FoxM1* mRNAs by QRT-PCR; C, Tubular injury scores. **D,** Representative images of PAS-stained sections showing areas with flattened epithelia (green arrow) and peritubular inflammatory cells (red arrow) in the OSOM of uninjured *PEPCK Cre+* kidneys. Results expressed as means +/- SEM, datapoints shown. T-tests compared results in *PEPCK Cre+* and *Cre-* mice, p values shown if <0.05. Scale bars=50μM.

To explore this further, we evaluated the functional characteristics of primary PTECs isolated from PTEC DN RAR and *Cre-* control mouse kidneys. Early passage (P2/3) cells used for our studies were enriched for PTECs with >80% expression of PTEC markers Kim-1 and AQP-1, and <1% aSMA^+^ myofibroblast contamination (S. Fig 8A). As anticipated, we found decreased expression of the RAR target genes, *RARb* and *Cy26B1*, and increased expression of *Kim-1* mRNA (and protein) in primary PTECs derived from PTEC DN RAR mice compared with *Cre-* controls (Fig. 7A/B, Fig. 8F/H). Cell counting also revealed that PTECs from PTEC DN RAR mice have more rapid growth rates in culture than *Cre-* controls (Fig. 7C). This is consistent with our data showing increased numbers of Ki67^+^ PTECs in PTEC DN-RAR mice after AKI.

**Figure 7.**
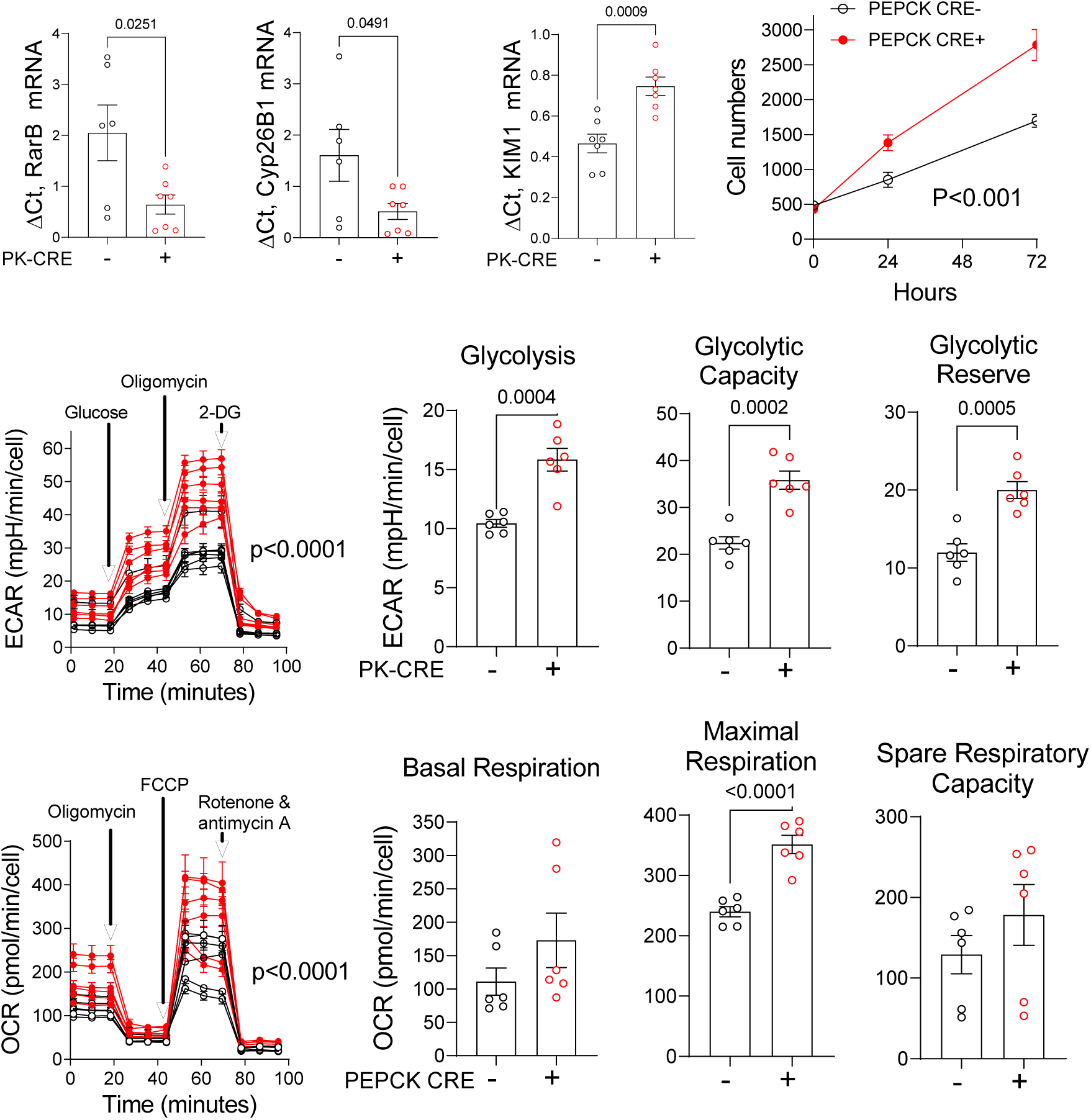
Inhibition of RAR signaling increases proliferation, glycolysis, and oxidative phosphorylation in cultured PTECs. Primary PTECs were isolated from a total of 7 uninjured *PEPCK Cre+* and 7 *Cre-* control mouse kidneys and evaluated at passages 2 and 3. Data are shown for biological replicates that were performed on separate preparations of primary PTECs isolated from different mice. **A,** RAR target genes *RarB* and *Cy26B1* mRNAs by QRT-PCR. **B,** Expression of *Kim-1* mRNA. Results expressed as mean +/- SEM, with individual data points shown. **C,** Cell growth. Cell numbers were counted over time. Results expressed as mean +/- SEM for PTECs isolated from 4 *PEPCK Cre+* and 4 *Cre-* control mouse kidneys. ***D/E, Increased glycolysis.* D,** Glycolytic stress tests on PTECs from 6 *PEPCK Cre+* (red) and 6 *Cre-* (black) mouse kidneys. **E,** Quantitative analysis of glycolytic rate, maximal glycolytic capacity, and glycolytic reserve. ***F/G, Increased maximal mitochondrial respiration.* F,** Mitochondrial stress tests on the same *PEPCK Cre+* and *Cre-* PTECs. **G,** Quantification of basal mitochondrial respiration, maximal mitochondrial respiration, and spare respiratory capacity. Results expressed as mean +/- SEM, with individual data points shown. A/D/F, T-tests were used to compare results in *PEPCK Cre+* and *Cre-* PTECs, p values shown if <0.05. B/C/E, 2-way ANOVA used to compare between groups over time, q values shown.

**Figure 8.**
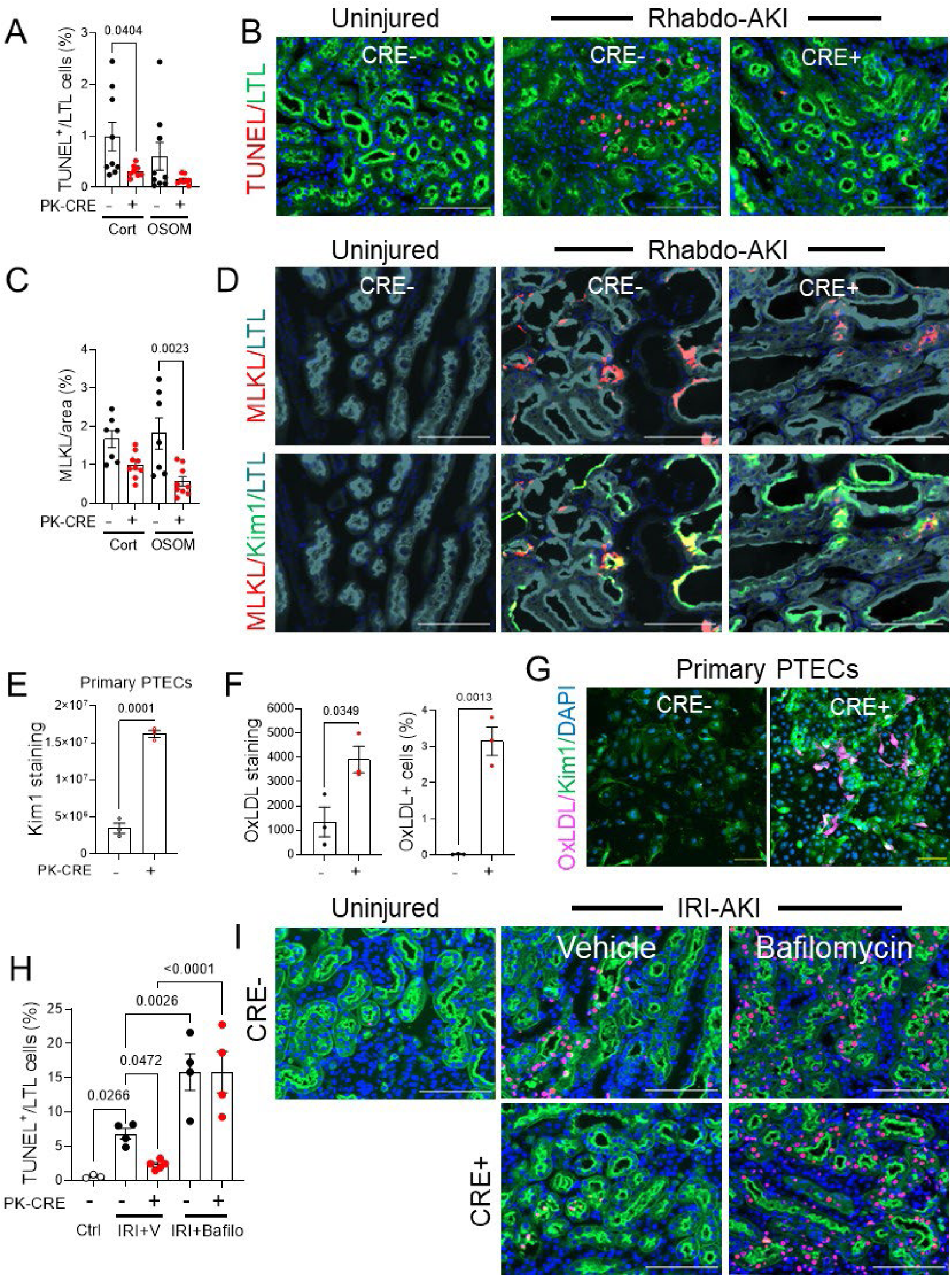
Increased efferocytosis in PTEC DN-RAR mice after AKI. *A-E, Decreased apoptosis in PTEC DN-RAR mice after rhabdo-AKI.* PTEC DN-RAR mice underwent rhabdo-AKI, and kidneys harvested after 3 days. **A,** Quantification of TUNEL+ LTL+ cells in the cortex and OSOM in *PEPCK Cre+* and *Cre-* controls. **B,** TUNEL and LTL staining in the OSOM in uninjured mice, and 3 days after rhabdo-AKI. **C,** Surface area staining with MLKL antibodies in the cortex and OSOM. **D,** Images showing MLKL with LTL, and LTL overlaid with Kim-1 immunostaining showing overlap of Kim-1 with MLKL in *PEPCK Cre-* kidneys. ***E-G, Kim-1 expression and endocytic activity.*** Primary PTECs were isolated from 3 *PEPCK Cre+* and 3 *Cre-* mice. **E,** Kim-1 fluorescence intensity. **F,** PTECs were incubated with fluorescently conjugated oxidized LDL (FC Ox-LDL), and cellular FC OxLDL evaluated after washing. Quantification of fluorescence intensity and numbers of PTECs taking up FC Ox-LDL. **G,** Kim-1 staining and FC Ox-LDL uptake in *PEPCK Cre+* and *Cre-* PTECs. ***H/I, Decreased apoptosis is due to increased lysosomal clearance of apoptotic cells in PTEC DN-RAR mice.* H,** PTEC DN-RAR mice were treated with bafilomycin A1, underwent bilateral IRI-AKI, and kidneys harvested after 24 hours. % TUNEL^+^ LTL^+^ cells in the OSOM. **I,** TUNEL and LTL staining. Results expressed as means +/- SEM, individual data points shown. A/C/F/G, T-tests, p values shown if <0.05. I, 1-way ANOVA, if p <0.05, q values shown for between group comparisons, corrected for repeated testing. B/D/E/J Scale bars=100mM, H, scale bars=20μM.

AKI is associated with mitochondrial damage giving rise to reduced mitochondrial oxidative metabolism, and increased glycolysis in de-differentiated, proliferating PTECs (61-63). Moreover, compared with other tubular segments, PTECs are thought to be particularly susceptible to injury because they have relatively low levels of glycolysis and are therefore dependent of mitochondria to generate ATP for solute transport functions and survival (61, 64). To explore the role of RAR signaling in these processes, we used Seahorse assays to compare cellular metabolism of primary PTECs isolated from PTEC DN RAR and *Cre-* control mice. PTECs from PTEC DN RAR mice have increased resting glycolysis, maximal glycolytic capacity, and glycolytic reserve (Fig. 7D/E), but also increased maximal respiratory capacity. There were no differences in basal respiration or spare respiratory capacity from mitochondrial oxidative metabolism compared with *Cre-* control PTECs (Fig. 7F/G). This was not associated with an increase in *Pgc1a* (which improves mitochondrial function (65)) or *Hk2* (required for glucose metabolism) mRNAs, but was associated with increased expression of *PfkB*, a rate limiting step for glycolysis (S. Fig. 8B) (62). These findings suggest that PTECs from PTEC DN RAR mice have metabolic features of de differentiated and regenerating PTECs (increased glycolysis), without evidence of injury-associated mitochondrial damage, since they have increased maximal respiratory capacity from mitochondrial oxidative metabolism. This also suggests that inhibition of RAR signaling induces metabolic reprogramming that could enhance the regenerative capacity of PTECs while at the same time protecting them from the damaging effects of ATP depletion in AKI.

### Increased Kim-1-dependent efferocytosis and suppression of inflammatory cell activation after AKI in PTEC DN RAR mice

Increased Kim-1 expression in PTEC DN RAR mice may reduce tubular injury by enhancing Kim-1-dependent apoptotic cell clearance, or efferocytosis (26-30). Consistent with this, there was a reduction in TUNEL^+^ PTECs (Fig. 8A/B), and a reduction in the expression of mixed lineage kinase domain-like protein (MLKL, a key necroptosis effector protein (66)) in PTEC DN RAR kidneys compared with *Cre-* controls after rhabdo-AKI (Fig. 8C/D). MLKL was membrane associated and largely co-localized with apical Kim-1 in *Cre-* control but not in *PEPCK Cre+* DN RAR mouse kidneys (Fig. 8D, S. Fig. 9). Since membrane-associated, apical MLKL is a terminal effector of necroptotic cell death in PTECs (67), these findings suggest that reduced PTEC apoptosis is also associated with reduced cellular necrosis. These findings would occur if there was increased Kim-1 dependent apoptotic cell clearance, or reduced apoptotic cell death. To distinguish between these two possibilities, we first sought to determine whether there was increased Kim-1 scavenging function in PTEC DN RAR mice.

In addition to its role as a phosphatidyl-serine receptor, Kim-1 mediates the endocytic uptake of oxidized lipoproteins that are also exposed on the surface of apoptotic cells (26, 29). Using primary PTECs isolated from PTEC DN RAR mice, which also have increased Kim-1 levels in culture, we found that there was increased cellular uptake of fluorescently conjugated oxidized LDL (Fig. 8E-G). This confirms that PTECs from PTEC DN RAR mice have increased Kim-1 functionality. We next evaluated whether blocking lysosomal degradation of apoptotic cells, which is required for Kim-1-dependent clearance of apoptotic cells (29), affects PTEC apoptosis in PTEC DN RAR mice after IRI-AKI. Like rhabdo-AKI, there was a reduction in TUNEL^+^ PTECs 24 hrs after bilateral IRI-AKI in PTEC DN RAR compared with *Cre-* control kidneys (Fig. 8H/I). To block lysosomal clearance of apoptotic cells, we treated mice with the vacuolar H^+^ ATPase inhibitor, bafilomycin A1, as described (29). By preventing lysosomal clearance of apoptotic cells that have been phagocytosed by PTECs via Kim-1 dependent efferocytosis, treatment with bafilomycin increased the number of apoptotic nuclei detected in PTECs after AKI. However, bafilomycin also reversed the effect of PTEC DN RAR mice on apoptotic PTECs 24 hrs after IRI-AKI (Fig. 8H/I). Since apoptotic cells are normally degraded and undetectable by TUNEL assay within 6 hours of being phagocytosed by PTECs (29), these findings indicate that reduced detection of apoptotic PTECs after AKI results from increased efferocytosis, rather than reduced PTEC apoptotic cell death, in PTEC DN RAR mice.

Apoptotic cell clearance by Kim-1 mediated efferocytosis suppresses the activation of macrophage-dependent inflammatory responses after AKI (29). We therefore asked whether there was a reduction in renal macrophage activation in PTEC DN RAR mice after AKI. Unexpectedly, we saw an increase in staining with F4/80 antibodies, a marker of tissue macrophages (68), in PTEC DN RAR mice after both rhabdo-AKI and IRI-AKI (Fig. 9A-D). F4/80 staining was also increased in uninjured PTEC DN RAR mouse kidneys (Fig. 9A/B). F4/80 staining was closely associated with Sox9^+^ PTECs in both uninjured PTEC DN RAR mice and injured mice after rhabdo-AKI (Fig. 9B). To explore this further, we performed RNA Seq on CD11B^+^ mononuclear cells isolated from PTEC DN RAR and *Cre-* control mouse kidneys 3 days after IRI-AKI. Principal component analysis (PCA) of these datasets showed clear separation of the two genotypes along the first dimension, accounting for 57% of the variation between samples (S. Fig. 10). Gene set enrichment analysis (GSEA) using the gene Ontology (GO) datasets demonstrated a marked downregulation of pro-inflammatory pathways in PTEC DN RAR vs. *Cre-* control CD11B^+^ cells in the kidney after IRI-AKI (Fig. 9E, S. Table 3). To evaluate this further, we performed GSEA using published datasets for validated pro-inflammatory (M1 activated) and anti-inflammatory (M2 activated) macrophage genes generated by transcriptional profiling of unstimulated mouse bone marrow derived macrophages (BMDMs), vs. BMDMs stimulated with LPS and g-IFN (to induce classical M1 activation) vs. IL4 (to induce anti-inflammatory, M2 macrophages) (S. Table 4 (69)). The same datasets have been used to characterize renal macrophage populations after AKI (70). There was a significant downregulation of genes that are uniquely upregulated in M1 activated macrophages (“distinct M1 up”), but also genes that are increased in both M1 and M2 macrophages compared with untreated control BMDMs (“increased M1+M2”), and in genes increased in M1 activated macrophages compared with untreated control BMDMs (“M1 up”) (Fig. 9E-G, S. Table 4). There was, however, no difference in expression of genes upregulated in M2 macrophages (“distinct M2 up”, and “M2 up”). These findings indicate that there is primarily a suppression of pro inflammatory macrophage activation markers in the kidneys of PTEC DN RAR mice after IRI-AKI.

**Figure 9.**
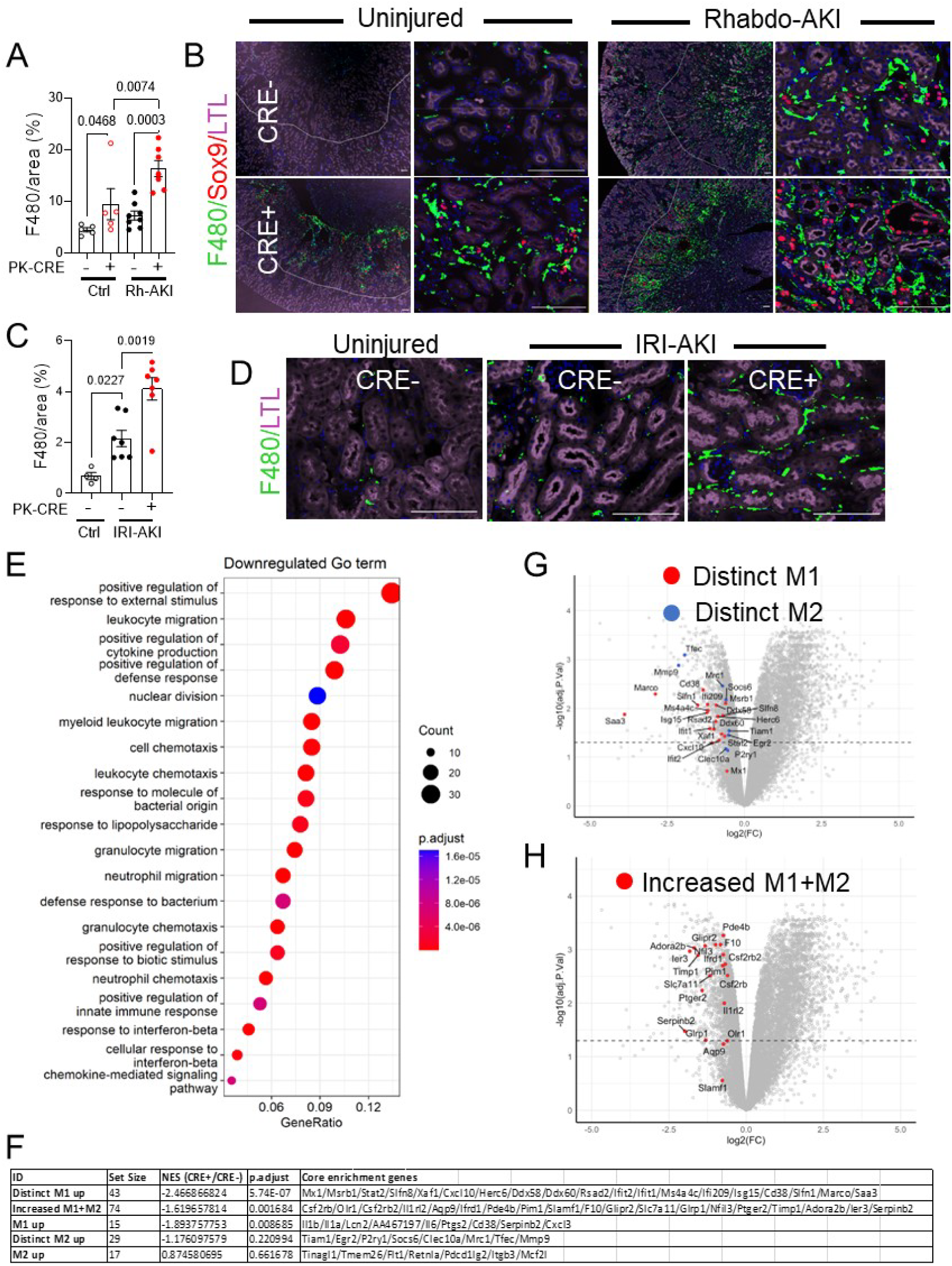
Increased renal macrophages with a reduced pro-inflammatory signature in PTEC DN-RAR mice after AKI. PTEC DN-RAR mice underwent rhabdo or bilateral IRI-AKI, and kidneys harvested after 3 days. ***A-D, Increased F4/80 staining in the kidney after rhabdo- and IRI-AKI.* A/B,** F4/80 staining in uninjured mice (Ctrl) and after rhabdo-AKI (Rh-AKI). Area staining with F4/80 in the OSOM, and representative images showing F4/80, Sox9, and LTL staining. Left hand panels show F4/80 staining is largely restricted to the OSOM. Right hand panels show higher magnification of the OSOM. **C/D,** F4/80 staining after bilateral IRI-AKI. Quantification, and images showing F4/80 and LTL staining in the OSOM. Results expressed as means +/- SEM, individual data points shown. A/C, 1-way ANOVA, if p <0.05, q values shown for between group comparisons. Scale bars=100μM. ***E-F, Decreased expression of inflammatory markers in CD11B+ cells after IRI-AKI.*** PTEC DN-RAR mice underwent bilateral IRI-AKI, and bulk RNA seq performed on renal CD11B^+^ cells 3 days after injury. **E,** Gene set enrichment analysis (GSEA) of downregulated genes using Gene Ontology (GO) datasets. **F,** GSEA for validated pro-inflammatory (“M1”), and anti-inflammatory (“M2”) gene sets in the RNA seq data. Set size=no. of genes from each gene set that are represented. NES=normalized expression score for the gene set sizes. **G/H,** Volcano plots showing fold change in expression of core enrichment gene from the CD11B+ bulk RNA seq dataset. Dotted line indicates p<0.05.

While CD11B^+^ cells are enriched for renal monocyte/macrophages, they include a number of other cell types including NK cells, granulocytes and B cells (71). To further define the phenotype of renal monocyte/macrophages in PTEC DN RAR mice, we performed flow cytometric analysis using a panel of mononuclear cell markers (S. Table 6E). CD45^+^ renal leukocytes were selected, while granulocytes (using Ly6G staining) and CD11B^-^ cells were excluded. Gating was then set to distinguish between F4/80^-^ (early infiltrating monocytes), F4/80^int^ (bone marrow-derived renal macrophages), and F4/80^hi^ (kidney resident macrophages, KRM) cells, as described (S. Fig 11) (70, 72). There was an expected increase in total CD45^+^, CD11B^+^, and CD45^+^; CD11B^+^ non-granulocyte (ly6G^-^) mononuclear cells after IRI-AKI, there was no difference in total cell numbers between *PEPCK Cre+* and *Cre-* kidneys (S. Fig 12). Consistent with our IF data, we found an increase in F4/80^hi^ cells in uninjured and IRI-AKI kidneys from PTEC DN RAR mice (Fig. 10A/B). KRM cells also express high levels of MHC Class II antigens (MHC-II^hi^) (70). As previously reported (70), we also detected high levels of MHC-II expression in F4/80^hi^ cells from uninjured mice, and there were fewer F4/80^hi^ MHC-II^hi^ cells after injury (S. Fig 13A/B). However, F4/80^hi^ MHC-II^hi^ cells were increased in PTEC DN RAR mice compared with *Cre-* controls after AKI. This confirms that the increase in F4/80^hi^ cells in PTEC DN RAR mice are KRMs. To determine whether the expansion of KRMs in PTEC DN RAR mice is due to increased proliferation, we performed pulse labeling with the S-phase marker, EdU. There was a significant increase in EdU^+^ F4/80^hi^ cells in PTEC DN RAR compared with *Cre-* controls after AKI (S. Fig 13C/D). This suggests that the increase numbers of F4/80^hi^ KRMs in PTEC DN RAR mouse kidneys arises, at least in part, from an increased rate of proliferation. We also noted a reduction in the cell inflammatory marker, Ly6C, in F4/80^-^ and F4/80^int^ infiltrating monocyte/macrophages, and bone marrow derived macrophages, respectively (Fig. 10C-F). Consistent with these findings, there was a reduction in cells staining with an anti-Gr1 antibody (which recognizes activated monocytes/macrophages <u>and</u> granulocytes (73)), but not MPO (which only recognizes granulocytes (74)) in PTEC DN RAR mice after IRI-AKI (S. Fig 13G-I). CD206/mannose receptor positive F4/80^hi^ cells, a marker of M2 activated macrophages (75), were also reduced in PTEC DN RAR kidneys after AKI, and there was a paradoxical increase in CD206^+^ F4/80^hi^ KRMs in uninjured PTEC DN RAR kidneys compared with *Cre-* controls (Fig. 10G/H). Expansion of M2 macrophages may explain why there is no evidence of tubular injury in uninjured PTEC DN RAR mouse kidneys despite the increase in F4/80^+^ KRMs.

**Figure 10.**
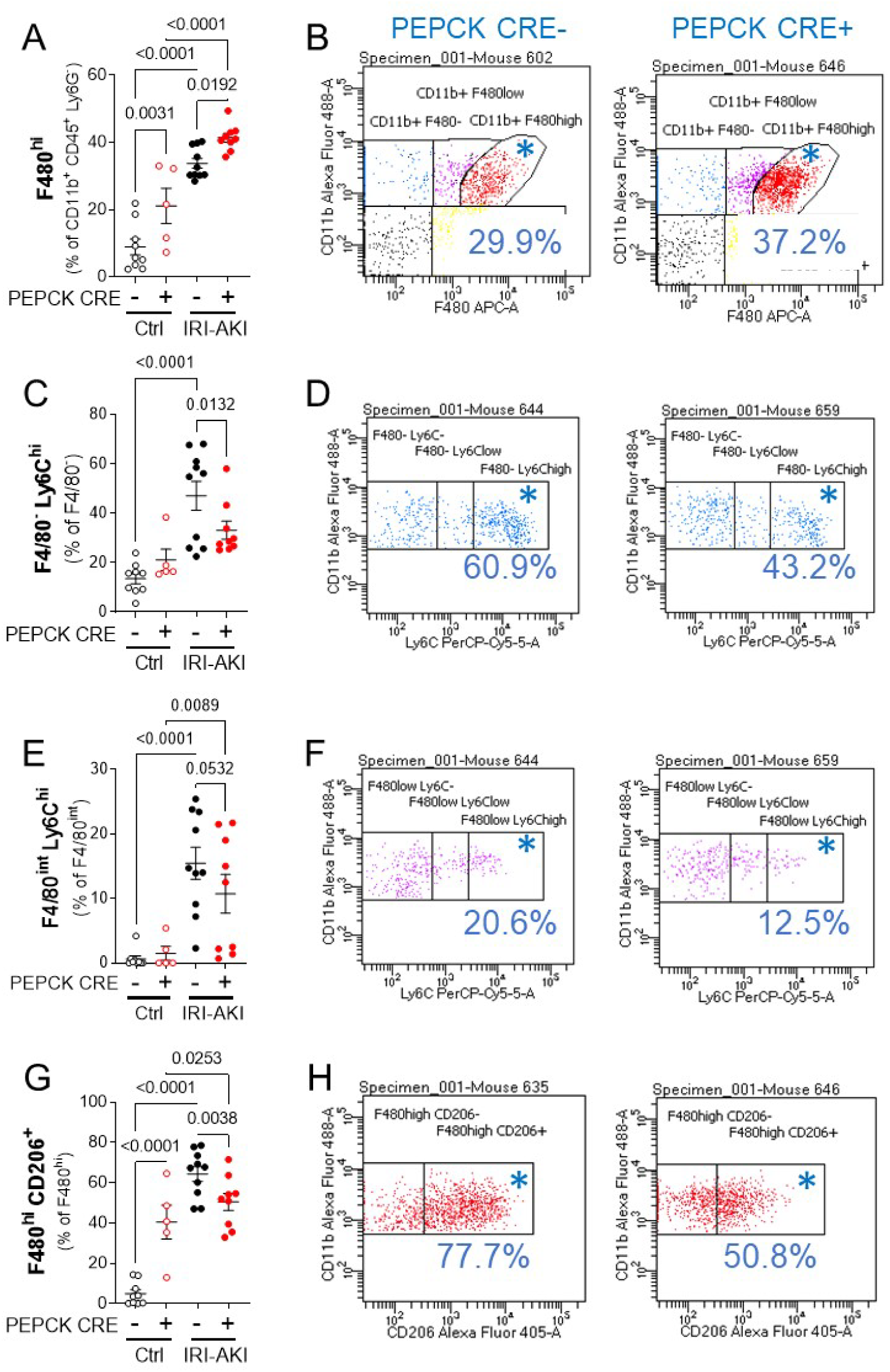
Flow cytometric analysis of renal monocyte/macrophages in PTEC DN RAR mice. Kidneys were harvested from PTEC DN RAR mice 3 days after bilateral IRI-AKI. Tissue was digested, homogenized, and evaluated by flow cytometry. ***A/B, F4/80^hi^ cells (kidney resident macrophages).*** A, F4/80^hi^ cells as the % of gated CD11B^+^ CD45^+^ Ly6G^-^ cells. **B,** CD11B and F4/80 expression charts in *PEPCK Cre+* and *Cre-* mice after IRI-AKI (% in gated area indicated with *). ***C-F, Ly6C^hi^ cells (inflammatory monocyte/macrophages)***. C, F4/80^-^ (infiltrating monocytes) Ly6C^hi^ cells, and E, F4/80^int^ (bone marrow derived macrophages) Ly6C^hi^ cells, as the % of gated F4/80^-^ and F4/80^int^ cells, respectively. D/F, CD11B and Ly6C expression charts. ***G/H, CD206/mannose receptor expression (an M2 activated macrophage marker)***. G, CD206^+^ cells as the % of gated F4/80^hi^ cells. H, CD11B and CD206 expression charts. Results are expressed as means +/- SEM, with individual datapoints shown. 1-way ANOVA was used to compare between groups, and if p <0.05, q values shown for between group comparisons, corrected for repeated testing.

These data indicate that despite the increase in F4/80^+^ KRMs in PTEC DN RAR mice, there is suppression of pro inflammatory monocyte/macrophage activation in the absence of injury and after AKI. Since Kim-1 dependent apoptotic cell clearance results in the suppression of early macrophage-dependent inflammatory responses after AKI (29), this is consistent with our findings of increased Kim-1 dependent clearance of apoptotic cells in PTEC DN RAR mice. In addition, since early pro-inflammatory macrophage responses worsen injury after rhabdo- and IRI-AKI (76-78), these findings may account for the reduction in severity of IRI-AKI and rhabdo-AKI after inhibition of RAR signaling in PTECs. This contrasts with an earlier study showing that long-term over-expression of Kim-1 in epithelial lineages increases expression of inflammatory markers, and increases renal fibrosis (33), and suggests that while short-term upregulation of Kim-1 in PTEC DN RAR mice protects against AKI, long-term Kim-1 over-expression results in pro-inflammatory kidney injury and fibrosis.

## Discussion

Our analysis of human RNA Seq data (11) shows that there is enrichment for RAR target genes in the kidneys of patients with severe SA-AKI, and that these are dominantly expressed by PTECs. The most highly enriched RAR target genes in SA AKI PTECs are also upregulated in mouse kidneys after rhabdo-AKI. Since RAR signaling was less extensively activated in PTECs after IRI than rhabdo-AKI (44), these findings also suggest that the mechanisms regulating RAR signaling in human SA-AKI maybe more closely related to rhabdo than IRI-AKI. It is notable that circulating cell free hemoglobin (CFH) detected in patients with sepsis can induce heme-dependent PTEC injury and exacerbate AKI mouse models of SA-AKI (79, 80). Since heme injury to PTECs is also the dominant mechanism of injury in rhabdo-AKI (81, 82), it is possible that CFH and myoglobin, both of which are reabsorbed from the glomerular filtrate in the proximal S1 and S2 PTEC segments in the renal cortex (83-86), explains the more widespread activation of RAR signaling in SA- and rhabdo-AKI compared to IRI-AKI, where PTEC injury and RAR activation is more restricted to S3 segment PTECs in the OSOM (14).

In addition to differences in the distribution of RAR signaling in rhabdo- and IRI-AKI, our studies show that inhibition of RAR signaling in PTECs is more protective in mice with rhabdo than IRI-AKI. Unlike IRI-AKI, there was significant improvement in renal function and reduction in tubular injury in both the cortex and OSOM in PTEC DN RAR mice after rhabdo-AKI. Because RAR activation extends throughout the cortex and outer medulla after rhabdo-AKI, but is more restricted to the outer medulla after IRI-AKI (44), this suggests that the net effect of inhibiting RAR signaling in PTECs on renal outcomes after AKI is affected by the differences in the extent and cellular distribution of RAR activation in the two models. In addition, while RAR signaling is dominantly activated in Kim-1^+^ PTECs after IRI-AKI, there was extensive RAR activation in Kim-1^-^ PTECs after rhabdo-AKI. Since Kim-1 is a marker of injured PTECs (20-23), this may be because RAR signaling is being activated in PTECs with less severe injury after rhabdo than after IRI-AKI. One consequence of this might be that Kim-1-mediated effects in PTEC DN RAR mice are more pronounced after rhabdo than IRI-AKI, since PTECs that would otherwise have been Kim-1^-^ are now induced to express Kim-1. This contrasts with IRI-AKI, where most of the PTECs with active RAR signaling are already Kim-1^+^ without inhibition of RAR signaling in PTEC DN RAR mice. These findings underscore the importance of using more than one model of human disease to understand relevance to cellular pathophysiologies in human AKI (13). In this case, while RAR signaling is activated in both experimental models of AKI, and since inhibition of PTEC RAR signaling exerts similar effects of PTEC function in both models, differences in the overall effects of inhibiting this pathway in PTECs depend on differences in the distribution of RAR signaling between models. Since there is also appears to be widespread activation of RAR signaling in PTECs from patients with SA-AKI, this also suggests that the effects of inhibiting PTEC RAR signaling in human SA-AKI may be better modeled by rhabdo than IRI-AKI.

Our studies provide the first *in vivo* evidence that Kim-1 up-regulation has protective effects in AKI. The protective effects of Kim-1 in promoting apoptotic cell clearance has been inferred from *in vitro* studies (26, 27), and from loss of function studies *in vivo* (28-30), but has not been evaluated *in vivo* by over-expression of Kim-1. In our studies, we show that the protective effects of inhibiting PTEC RAR signaling are associated with increased clearance of apoptotic cells, and a reduction in pro-inflammatory renal macrophages. Phagocytic clearance of apoptotic cells, including Kim-1 dependent efferocytosis by PTECs after AKI (29), reduces inflammation in different organ injuries (87, 88). This is mediated by reducing secondary necrosis resulting from the perseverance of apoptotic cells in injured tissues, and by direct anti inflammatory effects of efferocytosis on the phagocytic cell (87, 88). However, we also show that PTECs from PTEC DN RAR mice are de-differentiated, with increased Sox9 and FoxM1 expression, increased glycolysis, and increased mitochondrial oxidative phosphorylation. Given that increased Sox9 expression (89-91), and increased oxidative phosphorylation in PTECs (62, 63, 65), have protective effects on injured PTECs, it is possible that inhibition of RAR signaling in PTECs makes them more resilient to injury. In addition, our studies show that PTECs from DN RAR mice are more proliferative *in vitro* and *in vivo* after IRI- and rhabdo-AKI. This may be due to the increased rates of PTEC glycolysis, which would provide the metabolic building blocks for proliferative repair after injury (61), and suggests that surviving PTECs in PTEC DN RAR mice may have more robust regenerative responses than wild type PTECs with intact RAR signaling. However, if the dominant mechanism by which PTEC DN RAR mice are protected from AKI is mediated by direct cytoprotective effects on PTECs, and/or increased PTEC repair after injury, we would have anticipated: a) that reduced apoptosis in PTEC DN RAR mice would persist after inhibition of lysosomal degradation of apoptotic cells with bafilomycin; and/or b) long-term renal outcomes would improve as a result of increased regenerative repair. However, bafilomycin increased apoptotic PTECs in PTEC DN RAR mice with IRI-AKI, and there were no protective effects were seen on long term renal outcomes in PTEC DN RAR mice after IRI- and rhabdo-AKI. This suggests that protective effects of inhibiting PTEC DN RAR signaling require efferocytosis, and do not result from increased PTEC regeneration after AKI. These findings suggest that while additional ameliorating mechanisms maybe involved, the dominant mechanism by which inhibition of PTEC RAR signaling protects against AKI is by increasing Kim-1 dependent efferocytosis. These data also suggest a mechanisms by which persistent Kim-1 expression in PTECs that have failed to undergo productive repair after injury (92, 93), could confer a survival benefit to the affected individual by reducing further tubular injury after AKI.

In general, renal macrophages increase with worsening injury, and macrophage depletion studies indicate that early increases in renal macrophages worsen injury after AKI (76-78). Moreover, genetic studies have shown that loss of Kim-1 function, or expression increases renal macrophage numbers after AKI (29, 30). On this basis, we had anticipated that reduced renal injury in PTEC DN RAR mice would have been associated with reduced, not increased, renal macrophage numbers. Prior studies using a conditional genetic approach to induce widespread over-expression of Kim-1 in renal epithelium showed that persistent Kim-1 over-expression increased renal macrophages, expression of inflammatory markers, and caused progressive renal fibrosis (33). This has been used to support the hypothesis that persistent Kim-1 expression in PTECs promotes renal inflammation and fibrosis (33). However, this is also consistent with our finding that upregulation of Kim-1 both in injured and uninjured PTEC DN RAR mice is associated with increased numbers of renal macrophages. This suggests that Kim-1 over-expression alone may increase local recruitment and/or expansion of renal macrophages. This is supported by our observation that there is a close physical association between F4/80^+^ cells and Sox9^+^/Kim-1^+^ PTECs in PTEC DN RAR mouse kidneys, suggesting that these cells are recruiting and/or causing expansion of renal macrophages locally. The latter possibility is consistent with our finding that there are increased numbers of proliferating, EdU^+^ F4/80^Hi^ KRMs in PTEC DN RAR mice after injury. However, unlike the long-term, conditional Kim-1 over expression studies mentioned above (33), our data also shows that there is a suppression of inflammatory cell activation in both uninjured and injured PTEC DN RAR mouse kidneys after IRI-AKI. This may explain why renal macrophage recruitment by Kim-1 up-regulation in PTEC DN RAR mice does not promote renal injury.

Kim-1 is increased in PTEC DN RAR mice after AKI despite there being a reduction in tubular injury. Our studies show that this is associated with increased expression of Sox9, a transcription factor that is expressed by de-differentiated PTECs (59, 60), and Ki67, a marker of cell proliferation, in PTECs. In addition, the majority of Sox9^+^ PTECs are also Kim-1^+^ in both injured and uninjured PTEC DN RAR mouse kidneys. Cultured PTECs from PTEC DN RAR mice have increased expression of Kim-1, more rapid cell growth, and increased glycolytic flux, a metabolic feature of de-differentiated and proliferating PTECs (61-63). This is consistent with the observation that Kim-1 expression is localized to de-differentiated, proliferating PTECs after IRI-AKI (19), and in a number of models of toxin-induced AKI (20). This is also consistent with the finding that there is increased *Havcr1* in *Sox9*^+^ PTECs that have survived but have failed to undergo productive repair after IRI-AKI, and in patients with diabetic kidney disease (http://www.humphreyslab.com/SingleCell/) (92, 93). Further work will be required to determine how RAR signaling regulates PTEC differentiation and Kim-1 expression, however these findings suggest that increased Kim-1 expression may be driven by primarily by de-differentiation and not by injury per se.

Our finding that PTEC DN RAR mice are protected from AKI was unexpected, since we had previously shown that systemic inhibition of RAR signaling using the pan-RAR inhibitor, BMS493, worsened injury and increased long-term fibrosis after IRI-AKI (44). Moreover, systemic administration of RA reduces injury, decreases inflammatory cytokine expression, and reduces renal fibrosis after IRI-AKI (44). On this basis, we had anticipated that inhibition of RAR signaling in PTECs would have worsened, not ameliorated, AKI. However, we also showed that BMS493 decreases RAR activation in renal macrophages, and increases the expression of pro-inflammatory macrophage markers after IRI-AKI. In addition, worsening renal injury is reversed by depletion of macrophages, indicating that the detrimental effects of BMS493 in AKI are primarily dependent on macrophage-dependent injury, and not on the inhibitory effects of BMS493 on PTEC RAR signaling. These findings are consistent with the known anti-inflammatory effects on RA in cultured macrophages (47-49, 94), and suggest compartment specific effects of RAR signaling have opposing effects on renal outcomes after AKI.

There are a number of examples of signaling pathways that exert opposing effects on renal outcomes in renal macrophages vs. PTECs after kidney injury. For example, there is activation of the canonical mediator of Wnt signaling, β catenin in renal macrophages and PTECs after AKI, and while β-catenin depletion in PTECs worsens injury after AKI (95), depletion of β-catenin in macrophages decreases tubular injury and fibrosis after ureteric obstruction(96). Likewise, TGF-β type 2 receptor depletion in macrophages reduces post-injury fibrosis in IRI-AKI (97), while TGF-β type 2 receptor depletion in PTECs worsens tubular injury and fibrosis after aristolochic acid-induced renal injury (98). However, unlike the canonical Wnt and TGF-β signaling pathways, our data show that inhibition of canonical RAR signaling in PTECs protects against tubular injury after AKI. This suggests that activation of RAR signaling in PTECs may actually worsen tubular injury after AKI, and raises questions about the evolutionary benefit of activating PTEC RAR signaling after AKI.

While our studies indicate that the protective effect of inhibiting RAR signaling in PTECs are mediated by increased Kim-1 dependent efferocytosis, our studies also show that this is associated with increased de-differentiation, proliferation, and metabolic reprogramming of PTECs. This suggests that activation of RAR signaling in PTECs may actually be a compensatory response that acts to promote or maintain the mature differentiation state and quiescence of surviving PTECs after AKI. These findings are consistent with the known effects of RA in promoting cellular differentiation and quiescence in a variety of different contexts. These include the differentiation of hematopoietic stem cells, pancreatic and ovarian cancer cells, and non-malignant mammary epithelial cells (38), the differentiation and suppression of tumor proliferation in patients with promyelocytic leukemia, neuroblastoma, and early-stage breast cancers (99), and the differentiation and quiescence of glomerular podocytes after injury (39, 40). Widespread activation of RAR signaling in Kim-1^-^ PTECs after rhabdo-AKI suggests that this response could play a role in preventing PTEC de-differentiation in areas with relatively minor injury. We therefore propose that activation of RAR signaling in the kidney after AKI plays a role in preserving renal function, thus ensuring animal survival by reducing de-differentiation and thereby preventing further loss of function of surviving PTECs from regions of the kidney with less severe tubular damage after a renal insult.

## Methods

### RNA sequencing studies

Differentially expressed gene lists comparing human control and AKI kidney sample bulk and snRNA sequencing were obtained from S. Table S2 in Hinze et al. Genome Medicine 2022 (see S. Methods for details about patient and control samples) (52). For mouse renal CD11B cell studies, whole kidneys were harvested after cardiac perfusion with 0.9% saline, kidneys were finely minced in DMEM with 0.3mg/ml Liberase DL (Sigma/Roche) and homogenized at 37°C in MACS C-tubes using the gentle MACS dissociator (Miltenyi Biotec), digestion neutralized with PEB buffer (2 mM EDTA, 1% BSA in PBS), and cell suspension passed through 40μm cell strainer. Cells were then incubated with anti-mouse CD11b MACS beads (Miltenyi Biotech, 130-126-725) for 15 minutes, and CD11b+ cells separated by loading cells into MACS separator. The cell pellet was re-suspended in 350μl of buffer RLT (Qiagen), and RNA extracted with RNeasy Plus Mini Kits (Qiagen, 74134). After RNA quality control was confirmed with RIN scores >7.9, cDNA library preparation was performed using a polyA-selected library preparation kit by the Vanderbilt Technologies for Advanced Genomics (VANTAGE) core facility. Multiplexed sequencing was performed on an Illumina sequencer, with 50 BP single ended sequence reads at 30M reads/sample. Demultiplexed FASTQ RNA Seq reads, and alignment were performed as outlined in S. Methods. Gene set enrichment analyses (GSEA) were performed using the clusterProlifer R package (100). Enrichment score was calculated by comparing the log fold change in each gene in the gene set with all other expressed genes. This represented as the normalized to the gene set size for each gene set to generate normalized enrichment scores (NES). GSEA was performed for Gene Ontology (GO) classification from the MSigDB collections (101, 102), as well as validated macrophage and RA target gene sets from published data (53, 69, 70), as described in the text.

### Mouse studies

Mice were maintained on a 12:12-h light-dark cycle with free access to standard food and water. Euthanasia was performed by cervical dislocation after anesthesia with inhaled isoflurane at the end of each experiment, or at humane end points. Experiments were approved by the Vanderbilt Institutional Animal Care and Use Committee.

#### Mouse mutant lines and strains

Male BALB/c mice were purchased from Charles River (Wilmington, MA). *RARE-hsp68-LacZ* (on a CD-1 background) were purchased from Jackson labs (54). *PEPCK-Cre* mice on a mixed 129svJ; C57Bl/6 background were obtained from Volker Haase (56), and *Rosa26-LSL-RARaT403X (R26R-DN RAR)* on a C57Bl/6 background were obtained from Cathy Mendelsohn (36). Mice were genotyped and identified from ear punch biopsies using allele specific PCR primers, and copy number variant (CNV) genotyping as performed to identify homozygous R26R-DN RAR mice (see S. Methods for additional details about CNV genotyping. All of the genotyping primers are listed in S. Table 5A).

#### AKI models

For all injury models, body weight was monitored throughout the study. Blood was collected for the determination of serum BUN, and kidneys harvested analysis at the indicated time points. Numbers were randomly assigned to each mouse, and split equally into groups for each study. Another individual performed assays and data analyses and was blinded to the group assignments until analyses were completed. Rhabdomyolysis-AKI (rhabdo-AKI) was induced in male mice aged 13-14wk were given intramuscular (IM) 50% glycerol (Invitrogen, Carlsbad, CA) in sterile water after water deprivation for 18hrs prior to injection, as described (55). Mice were euthanized on Day 3 or 14 after glycerol injection depending on the study. Unilateral IRI-AKI with delayed nephrectomy (IRI-AKI DN) was performed on 10–12 week old male PTEC DN RAR mice, as described (103). Transdermal glomerular filtration rates (tGFR) were measured at Day 27, as described (58), and mice euthanized at Day 28 after the initial surgery. Mice that underwent right nephrectomy alone were used as controls. For bilateral IRI-AKI studies, 10–12-week-old male PTEC DN RAR mice underwent bilateral renal pedicle clamping for 28 minutes, and were euthanized Day 1 or 3 after surgery depending on the study. For bafilomycin A1 studies, mice were injected IP with 3 mg/kg bafilomycin A1 (Med Chem Express LLC, 50-196-7747) in corn oil 1 hour before bilateral IRI-AKI surgery, as described (29). Control animals were administered the same amount of corn oil (see S. Methods for additional technical details about the IRI- and rhabdo-AKI models).

#### Assessment of renal function

Blood was collected by submandibular vein or cardiac puncture into lithium-heparin-coated microcuvette tubes. Plasma was collected for BUN and measured in duplicate, according to the manufacturer’s instructions (Infinity Urea, Thermo Scientific, Waltham, MA). Transdermal glomerular filtration rates (tGFR) was performed in conscious mice, as described (58). The FITC-sinistrin half-life was calculated using a three-compartment model with linear fit using MPD Studio software (MediBeacon, Mannheim, Germany). The FITC sinistrin half-life was converted to tGFR (in μl/min) with correction for mouse body weight, as described (58).

Tissue Analyses. Kidneys were collected after terminal cardiac perfusion for snap freezing (for RNA), and preparation of formalin fixed, frozen (FFF) and formalin fixed, paraffin embedded (FFPE) tissue blocks (see S. Methods for details).

#### β-Galactosidase and immunofluorescence staining and analysis

β-Galactosidase staining, and antibody co-labeling was performed on FFF sections, as described (44). Immunofluorescence (IF) studies were performed on FFPE or FFF sections depending on the antibodies, as outlined in S. Table 5B. Blocking and antibody incubation steps were performed using the universal blocking reagent (Biogenex, Tremont, CA). Lectins, primary and secondary antibodies, dilutions, biotin amplification, and respective antigen retrievalmethods are summarized in S.Table 5B/C. TUNEL staining was performed using TMR-Red in situ cell death detection kit (Roche, 12156792910) per the manufacturer’s instructions, and counterstained with LTL. For quantification of IF and β-galactosidase-stained images, digital images were scanned using Zeiss AxioScan Z1 slide scanner (Carl Zeiss Microscopy GmbH, Oberkochen, Germany, 10X), and downloaded into QuPath (version 0.4.3) to generate color overlays. For quantification, kidney regions were identified on whole kidney scanned images using established landmarks depending on the experiment. Images were then quantified either as surface area stained in the indicated regions, or the ratio of cells staining with the indicated markers using DAPI staining to quantify individual cell nuclei using QuPath. Cell numbers and cell types (e.g.: LTL^+^, and LacZ^+^ cells) in different kidney areas (i.e.: cortex, OSOM, or ISOM) were identified based on average values of fluorescence staining thresholding detection using machine learning classifiers in QuPath (see S. Methods for additional details).

#### Histological scoring

Renal tubular injury scores and fibrosis/collagen deposition were determined on FFPE sections after staining with Periodic Acid Schiff (PAS) and Sirius red (SR), respectively, by blinded observers, as described (55, 104). H.Y. evaluated PAS-stained sections to determine tubular injury scores (TIS), as described (55). R.D. evaluated SR staining, using an Olympus BX-41 microscope equipped with a polarized light filter, using ImageJ to quantify birefringent SR stained collagen fibrils areas/total surface areas from digitally captured images, as described (104).

#### RNA extraction and quantitative RT-PCR

RNA purification from tissues and cultured cells was performed using RNeasy Mini Kits, according to the manufacturer’s instructions (Qiagen, 74106). Frozen tissue was homogenized in the RNeasy lysis buffer mixed with ceramic beads (Lysing Matrix A tubes, MP Biomedicals 6910050) using the Mini Bead Beater (BioSpec Products Inc, 607). RNA was quantified using a Nanodrop Spectrophotometer, and cDNA synthesized from 1μg RNA with iScript cDNA synthesis kits (Bio-Rad 1708891). Quantitative PCR was performed using iQ SYBR Green super mix (Bio-Rad) using a Bio-Rad CFX96 real time PCR system. mRNA expression was normalized to *Gapdh* mRNA using CFX Maestro software, results reported as normalized expression (ΔΔCt). Primers pairs were downloaded directly from PrimerBank, or designed using the National Center for Biotechnology Information Primer Designing tool, as described (104). PrimerBlast was used to confirm that primers spanned exon junctions. Primer sequences are listed S. Table 5D.

### Primary proximal tubular epithelial cell culture

#### Isolation and characterization

Primary mouse PTECs were isolated, as described (105). Briefly, the kidney cortex was minced into small pieces, digested with collagenase (3 mg/mL, Worthington, S3N6800)/Dispase (1 mg/mL, Gibco, 17105-041)/DNAase (0.1 mg/mL, Sigma-Aldrich, 11284932001) in PBS, and passed through a 40μm strainer (Thermo Fisher Scientific, 22363547). PTECs labeled with biotinylated LTL (Vector Laboratories, B-1325-2) were separated on anti-biotin microbeads (Miltenyi Biotec, 130-090-485) and seeded on collagen 1-coated dishes (30μg/mL) and grown in DMEM/F12 media supplemented with 50 ng/mL hydrocortisone, 5μg/mL insulin/transferrin/selenium, 6.5 ng/mL triiodothyronine, murine EGF (20 ng/mL, Peprotech, 315-09) penicillin/ streptomycin, and 0.5% BSA (Sigma-Aldrich, A3059). Cells were used between passages 2 and 3. For IF staining, cells were plated uniformly at a density of 1 × 10^5^cells/well in an 8-well chamber slide (ThermoFisher, 154453 or iBidi, 80826). Cells were washed after reaching confluence, fixed with 5% Formalin and permeabilized with 0.1% Triton X-100 in PBS. After blocking with 2.5% BSA, cells were incubated with primary antibodies overnight at 4°C, and incubated with secondary antibodies for 1 hour. Primary and secondary antibodies used to characterize primary PTECs are summarized in S.Table 5B/C.

#### Seahorse assays

Extracellular acidification rates (ECAR, XF Glycolytic Stress Test) and oxygen consumption rates (OCR, XF Cell Mito Stress Test) were performed with a Seahorse XF24 Extracellular Flux Analyzer and test kits (Agilent Technologies, Inc., Santa Clara, CA, USA). Passage 2 PTECs were seeded overnight in quintuplicate at a density of 5 × 10^4^ cells per well on a Seahorse cell culture plate in PTEC complete media. For ECAR assays, PTECs were incubated in Seahorse assay medium (Agilent Technologies) supplemented with 1 mM L-glutamine in a 37°C incubator without CO2 for 45 min prior to starting the assay. Glucose, oligomycin and 2-DG were injected, according to the manufacturer’s instructions. For OCR assays, cells were washed and incubated in Seahorse assay medium supplemented with 1 mM sodium pyruvate, 2 mM L-glutamine, and 25 mM Glucose in a 37°C incubator without CO2 for 45 min. Oligomycin FCCP and rotenone/antimycin were injected per the manufacturer’s instructions. The ECAR (mpH/min) and OCR (pMoles O2/min) was measured in real time, and glycolytic and mitochondrial parameters calculated using Wave software (Agilent). After the assays, cells were fixed with 100% Methanol and stained with DAPI solution. Values were normalized for count of DAPI stained nuclei in each well.

#### Endocytosis of oxidized LDL

Cells were plated onto 8-well chamber slides (iBidi, 80826) at a density of 1 x 10^5^cells/well. After the cells became approximately 70 % confluent, medium was changed to serum-free DMEM media to starve the cells for 6 hours. Dil-oxLDL (20μg/ml, ThermoFisher, L34358) was added into the media for 2 hours incubated at 37°C. Cells were then washed 3 times, fixed with 5% Formalin-PBS and co-staining with Kim-1 antibody.

### Flow cytometry

Kidneys were homogenized to single cell suspensions, as outlined above, and residual RBCs lysed with RBC lysis buffer (eBioscience) before proceeding with staining. For dead cells exclusion, Live/Dead Fixable Dead Cell Stain was added to cells. Using anti-mouse CD16/32 to block nonspecific Fc binding, cells were then incubated with cell surface antibodies in FACS buffer for 30 minutes on ice in the dark. For intracellular (IC) antibody (CD206) staining, cells were fixed and permeabilized with fixation and permeabilization buffer(eBioscience) subsequently, then incubated with IC antibody the same as antibody above. For EdU detection, 1mg/mouse EdU (Invitrogen, C10424) was administrated IP 2- or 24- hours before sacrifice. Following cell surface antibody staining and cell fixation and permeabilization, cells were resuspended in Click-iT Plus reaction cocktail solution and incubated for 30min on ice. Additionally, fluorescence-minus one (FMO) controls were prepared for each cell marker. Data were collected on a LSR II (BD, San Jose, USA) equipped with five aligned 355, 405, 488,532 and 633 nm lasers. The data analysis including sequential gating based on FMO control was performed by using BD FACS diva software. All antibodies and fluorophores used are listed in S. Table 5E.

**Statistical analyses** were performed using GraphPad Prism 9.3 software (San Diego, CA). Two-tailed, unpaired T-Tests were used to compare two groups, 1-way ANOVA used to compare between 3 or more groups. If p values were <0.05 for between group analyses, multiple between group comparisons were performed controlling for false discovery rates (FDR) of p<0.05 using the two-step method of Benjamini, Krieger and Yekutieli. 2-way ANOVA was used to compare between two group over time. If p values were <0.05 for between group analyses, between group comparisons at each time point were performed controlling for false discovery rates (FDR) at p<0.05. Q values after are indicated on the graphs for ANOVA tests after correction for multiple between group comparisons. Mantel-Cox log-rank test was used to compare survival between *PEPCK-Cre+* and *Cre* groups. Mouse numbers for each study are indicated in the figures and figure legends. Note that because of the large numbers of mice in the PTEC DN-RAR studies, we randomly selected subsets of samples for analysis of histology, IF, and RNA studies before performing the assays, as indicated in the figures.

### Data availability

All relevant data can be found within this article and in the supplemental data. CD11B cell bulk RNA sequencing data has been deposited with the NCBIs Gene Expression Omnibus repository (give GSE number). Human RNA sequencing data used for GSEA analysis was obtained from differentially expressed gene lists in S. Table 2 and UMAP plots of the snRNA sequencing datasets (https://shiny.mdc-berlin.de/humAKI/) from Hinze et al. Genome Medicine 2022 (11).

## Author contributions

M.Y. designed, conducted, and analyzed experiments with assistance from L.L. M.B., K.C., Y.Z, R.D., A.M. helped conduct and analyze experiments. H.Y. performed tubular injury scoring. T.V. and A.D. assisted with data analysis (GSEA studies). R.H., L.G., and C.B., helped design studies and interpret data, R.H., L.G., C.B. and A.D., edited the manuscript and figures. M.D.C. conceptualized and planned studies, interpreted data, prepared figures, and wrote the manuscript.

## Acknowledgements

Funding for this work: R01DK112688 and DODPR161028 (M.D.C), UC2DK126122 (A.D., M.D.C), 5RO1DK95785 (R.H.), RO1DK121101 (C.B), and RO1DK108968 (L.G.). RNA sequencing was performed by the Vanderbilt Technologies for Advanced Genomics core. Swapna Menon, Bioinformatics Specialist, Analyze Dat Consulting Services, Kerala, India, performed alignment and gene annotations on the mouse RNA seq dataset. The graphical abstract was created using BioRender.com.

## Supplemental Methods

### Validation of the human AKI RNA sequencing dataset

Differentially expressed gene lists comparing control and AKI kidney samples evaluated by bulk and snRNA sequencing for different renal cells data were obtained from S. Table S2 in Hinze et al. Genome Medicine 2022.^1^ RNA and snRNA samples were obtained from cadaveric renal biopsies obtained 60-120 minutes after cessation of circulation in 8 patients who died in a clinical care setting with pneumonia associated with severe sepsis-associated AKI (KDIGO 2 and 3) diagnosed within 5 days of sampling . Three age matched control samples were obtained from tumor adjacent normal renal tissue obtained at nephrectomy, and one control for validation of cadaveric donor renal biopsies obtained from timed post mortem biopsies (15, 60 and 120 minutes after circulatory arrest) in a younger patient with brain death. Principal component analysis (PCA) showed clear separation of control and AKI samples, and control samples obtained after 15, 60 and 120 minutes of circulatory arrest clustered closely with nephrectomy specimens in the two major dimensions (Dm1 and Dm2, accounting for 38 and 13% of variance, respectively). snRNA seq analysis demonstrated very similar distributions of the 12 identified renal cell types between all control and AKI samples. Note that PTECs and TAL cells accounted for ∼60% of the cells identified from snRNA seq analysis, with lower %’s of CD and leukocyte populations (∼10% and 2% respectively).

### Mouse RNA sequencing alignment and analysis

Demultiplexed FASTQ RNA Seq reads were aligned using STAR 2.5.0a, controlling for unannotated, non-canonical and rare junctions. Mouse genome assembly, GRCm38, and corresponding transcript annotations, downloaded from Ensembl, were used for alignment. The htseq-count function from HTSeq 0.6.1 was used in the intersection, non-empty mode to count the aligned reads. The edgeR and limma packages were used for differential expression analysis between the two conditions. ^2^ ^3^ Genes with log fold change ≥ 1 or ≤ -1 and adjusted p value < 0.05 are considered as differentially expressed.

### Mouse mutant lines

Note: *RARE-hsp68-LacZ* were also backcrossed for 9 generations onto a BALB/c background, but *RARE-LacZ* expression was no longer detectable in the kidney after rhabdo-AKI, so this background strain was discontinued. To select for homozygous *R26R-DN RAR* mice, we performed copy number variant (CNV) genotyping. For this, DNA was extracted using DNeasy Blood and Tissue Mini Kits, per the manufacturer’s instructions (Qiagen, 69504). DNA was quantified using a Nanodrop Spectrophotometer, and quantitative PCR performed in triplicate using 10ng of sample DNA with iQ SYBR Green super mix (Bio-Rad) using a Bio-Rad CFX96 real time PCR system. Known wild type, heterozygous and homozygous *R26R DN RAR* DNA samples (established from breeding) were included in each experiment as controls. DNA copy number was determined with primers for *Gapdh* as the reference genomic DNA control using CFX Maestro software: wild type mice have *R26R DN RAR* copy numbers close to 0, heterozygous mice between 0.4 and 0.6, and homozygous mice 0.8 to 1.2. Primer sequences used for CNV genotyping are listed in S. Table 5A.

### AKI models

To induce rhabdo-AKI, wild type BALB/c mice were given 6ml/kg glycerol; *RARE-LacZ* mice, 5.5ml/kg glycerol; and PTEC DN RAR mice 5.8ml/kg glycerol. Glycerol was injected into anterior thigh muscles while mice were anesthetized with a ketamine-xylazine mixture (120–150 mg/kg ketamine and 12–15 mg/kg xylazine) IP, with the glycerol dose split equally between the two hindlimbs. Pain relief with Buprenorphine was provided for 48 h starting immediately after intramuscular injections. Mice were also given 0.5ml of subcutaneous (SC) sterile 0.9% saline immediately after and 24 h after intramuscular injections. For bilateral IRI-AKI, 10–12-week-old male PTEC DN RAR mice were anaesthetized with IP ketamine-xylazine mixture and placed prone on a surface heated by a circulating water bath at 38°C. The kidneys were exteriorized via a dorsal incision, and after dissection of the renal pedicle from perinephric fat, placed back into the retroperitoneal space for 8 minutes to allow the kidneys to return to body temperature. 200-240gm pressure clamps (Roboz RS-5459) clamps were then placed on both renal pedicles staggered 1-2 minutes apart, and kidneys returned to the retroperitoneal space, and the mice was covered with a dry gauze pad and foil blanket. After 28 minutes, kidneys were exteriorized, the clamps removed, and kidney observed over 60 seconds for return of pink coloration before returning them to the retroperitoneal space and closing the wound in layers. If kidneys did not fully re-perfuse or there were other technical problems with the surgery, mice were excluded from analyses. 0.5 ml of 0.9% saline was given SC immediately after and the morning after surgery. Pain relief was provided with buprenorphine for 48hrs. All surgeries were started in the morning and completed by noon. For unilateral IRI-AKI with delayed nephrectomy, mice were anaesthetized with a ketamine-xylazine mixture, the left kidney exteriorized, the renal pedicle clamped and then removed after a defined time period . Studies were performed in two separate experiments with 34- and 36-minutes renal pedicle clamp times with no differences in survival or renal function between studies. For the nephrectomy surgery, mice were anesthetized using inhaled isoflurane, the right kidney exteriorized, the renal pedicle was tied off with silk suture, and the kidney removed. Pain relief was provided with SC buprenorphine for 48 h after surgery. Surgeries were started in the morning and completed by noon, and 0.5 ml of SC 0.9% saline was given immediately after and the morning after IRI surgery.

### Tissue harvesting and immunofluorescence staining

Mice underwent terminal cardiac perfusion under anesthesia, initially with 0.9% saline solution and right kidney clamped and removed. 2-3mm transverse blocks were cut transversely through the central cortex and medulla and allocated for formalin fixation and paraffin embedding (FFPE) after immersion in 10% buffered formalin (BF, Fisher brand) for 4 hours, or blocks were snap frozen in liquid nitrogen for subsequent RNA extraction. Mice then continued with cardiac perfusion with 10% BF and the left kidney removed and fixed in 10% BF for another hour. After cutting transverse blocks and washing, samples were placed in 30% sucrose in phosphate buffered saline (PBS) overnight, mounted in Tissue-Tek OCT (Sakura), and stored at -80C for subsequent immunofluorescence or b Galactosidase staining assays. For co-labeling studies after b-Galactosidase staining, after incubation with X-Gal substrate, sections were fixed in methanol, before antigen retrieval, blocking steps and incubation with primary and secondary antibodies, and/or biotinylated lectins. TUNEL staining was performed using TMR-Red in situ cell death detection kit (Roche, 12156792910) according to the manufacturer’s instructions. Briefly, after FFPE slides were de-paraffinized, sections were incubated with Proteinase K (20µg/ml) for 10 min at 37 °C. Slides were then incubated with TUNEL reaction mixture (5µl of Enzyme solution, 45µl of Label solution) at 37 °C for 1 hour, followed by washing in PBS.

### Image analysis

For quantification of IF and b-galactosidase-stained images, digital images were scanned using Zeiss AxioScan Z1 slide scanner with EGFP, DAPI, DsRed, and AF 647 filters on (Carl Zeiss Microscopy GmbH, Oberkochen, Germany, 10X), and digital images downloaded into QuPath (version 0.4.3) to generate color overlays. For b-Gal/ IF staining overlays, the bright field blue X-Gal color change was pseudo colored in white and overlaid onto digitally acquired fluorescence images. For quantification, kidney regions (cortex, outer stripe of the outer medulla (OSOM), inner stripe of the outer medulla (ISOM) and inner medulla (IM)) were identified on whole kidney scanned images using established landmarks depending on the experiment (e.g.: juxta medullary glomeruli to demarcate the cortex/OSOM junction, LTL staining the OSOM/ISOM junction; and THP-1 staining to demarcate the ISOM/IM junction). Images were then quantified either as surface area stained in the indicated regions, or the ratio of cells staining with the indicated markers using DAPI staining to quantify individual cell nuclei, as indicated in the figure legends, using QuPath. Cell numbers and cell types (e.g.: LTL^+^, and LacZ^+^ cells) in different kidney areas (i.e.: cortex, OSOM, or ISOM), were identified based on average values of fluorescence staining using machine learning and thresholding detection using QuPath. Briefly, the pixel classifier was trained with a minimum of 30 training regions for the cell (DAPI stained nuclei) and cell type (e.g.: LTL^+^ tubules), and identified as areas of fluorescence above background levels (nucleus threshold, segmentation parameters, cell expansion, smoothed features). Once the machine learning classifier was created, LTL^+^ (or other marker) were detected in the entire tissue section. Positive cell detection was then used to identify the labeled proteins (e.g.: LacZ/ Sox9) by thresholding, and these determined in cell types (e.g.: LTL^+^). Specific protein expression localization with respect to the LTL^+^ tubules was analyzed.

## Supplemental Tables

**S. Table 1. RAR target genes lists used for GSEA of human RNA Seq data. A,** Validated RAR target genes from Balmer, et al. 2002 ^4^. **B,** RA signaling regulators previously shown to be upregulated by RAR signaling after IRI-AKI 5.

**S. Table 2. GSEA analysis of RAR target genes in human SA-AKI RNA Seq databases from Hinze, et al. 2022^6^.** Tab labels indicate GSEA analysis on bulk RNA Seq data for the whole kidney, PTECs, leukocytes, TAL, collecting duct principal cells, interstitial cells and endothelial cells.

**S. Table 3. GSEA for the GO in mouse kidney CD11B bulk RNA Seq data from PTEC DN RAR mice after IRI-AKI.** Tab labels indicate summary of the analysis, GO term upregulated and downregulated gene sets along with core enrichment gene IDs driving significant associations with the bulk RNA Seq data.

**S. Table 4. M1 and M2 activated macrophage gene lists used for GSEA analysis CD11B bulk RNA Seq data.** Validated macrophage marked gene set obtained from Jablonski, et al. 2015 ^7^. Tab labels indicate individual gene sets.

**S. Table 5. Primers and antibodies used for genotyping, QRT-PCR, immunofluorescence, and flow cytometry studies. A,** Genotyping primer pairs. **B,** Primary antibodies and lectins used for immunofluorescence studies. **C,** Secondary antibodies and fluorophores used from immunofluorescence studies. **D,** Primers used for QRT-PCR studies. **E,** Antibodies and fluorophores used for flow cytometry studies.

---

**S. Figure 1.**
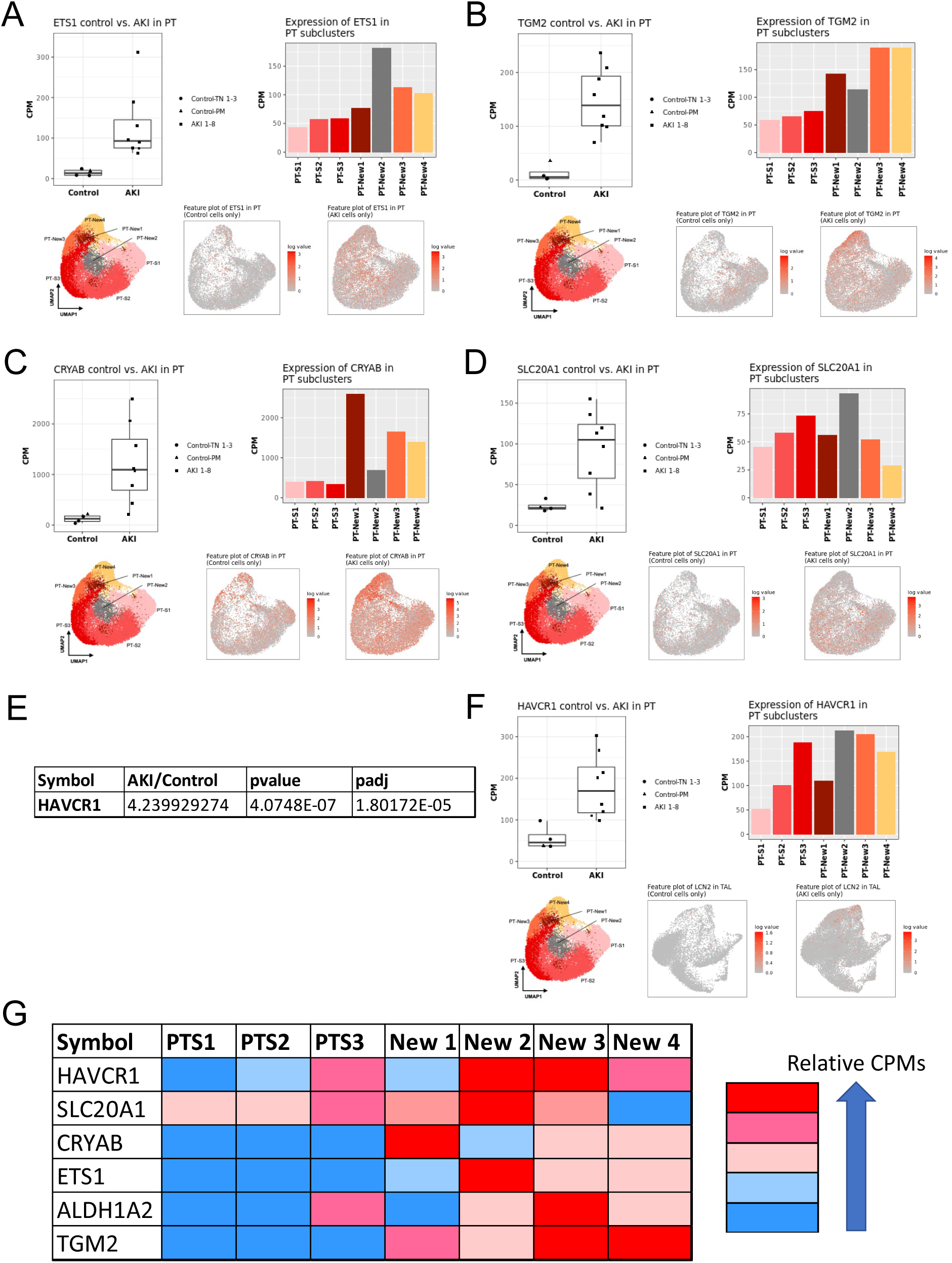
Expression of RAR target genes in proximal tubular epithelial cells from patients with SA-AKI. **A-D,** Expression of RAR core enrichment genes identified from GSEA analysis of the PTEC snRNA seq dataset from Hinze et al. 2022. **E/F,** Expression of Kim-1/Havcr1. Figures were generated using on-line software developed for these datasets at https://shiny.mdc-berlin.de/humAKI/. Gene expression in controls vs. AKI patients in reads per million. UPMAP plots indicate PTEC subclusters identified from snRNA seq studies. Feature plots showing distribution of genes in PTECs clusters summarized in bar charts. G, Visual representation of the distribution of RAR target genes in PTEC clusters.

**S. Figure 2.**
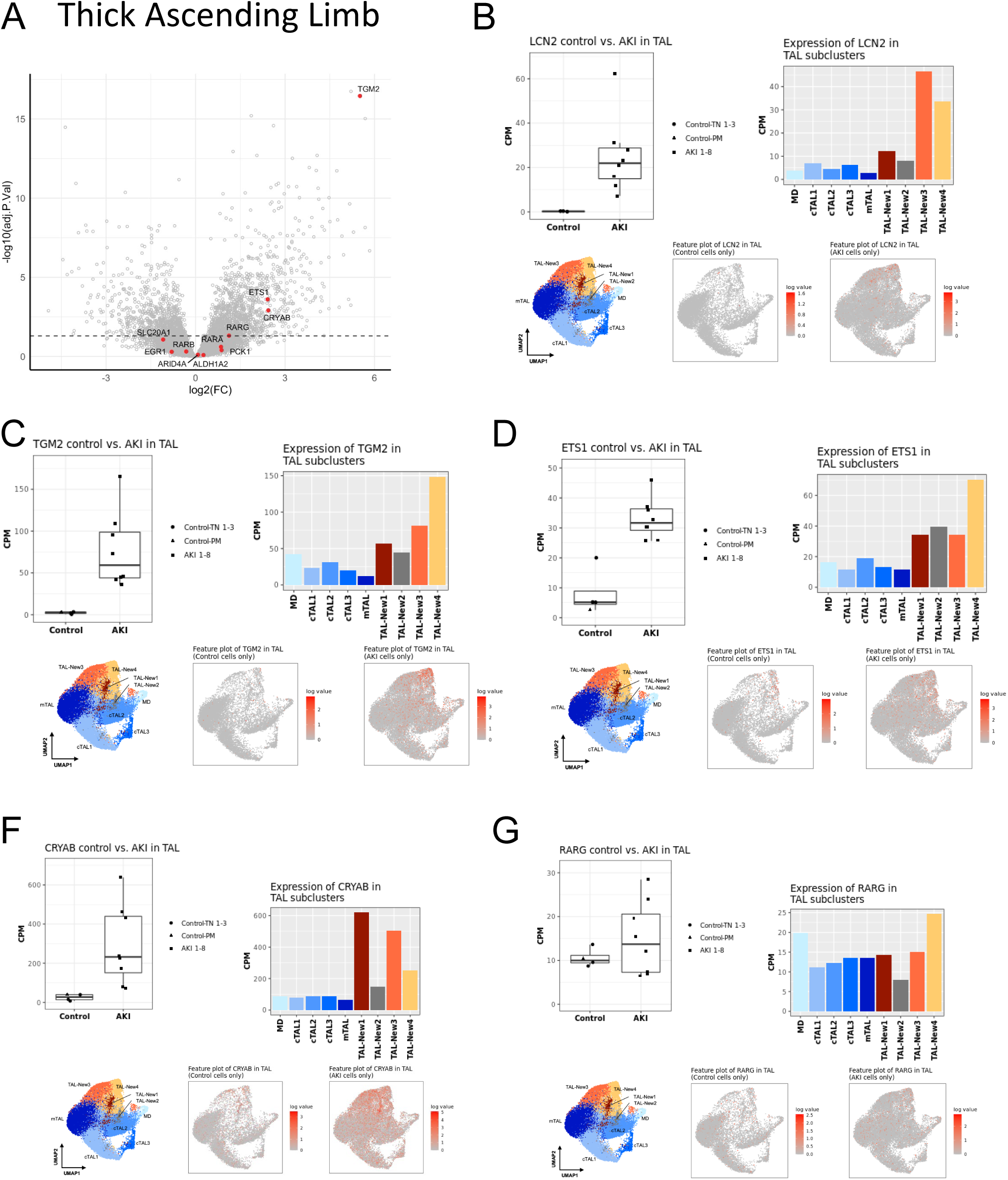
Expression of RAR target genes in thick ascending limb cells from patients with SA-AKI. **A,** Volcano plot indicating fold change in expression of core enrichment gene (AKI vs. controls) from the TAL snRNA seq dataset from Hinze et al. 2022. Dotted line indicates p<0.05E. **B,** Expression of N-Gal/Lcn2. **C-G,** Expression of RAR core enrichment genes identified from GSEA analysis of the TAL snRNA seq dataset from Hinze et al. 2022.^1^ Figures were generated using on-line software developed for these datasets at https://shiny.mdc-berlin.de/humAKI/. Gene expression in controls vs. AKI patients in reads per million. UPMAP plots indicate PTEC subclusters identified from snRNA seq studies. Feature plots showing distribution of genes in TAL subclusters summarized in bar charts.

**S. Figure 3.**
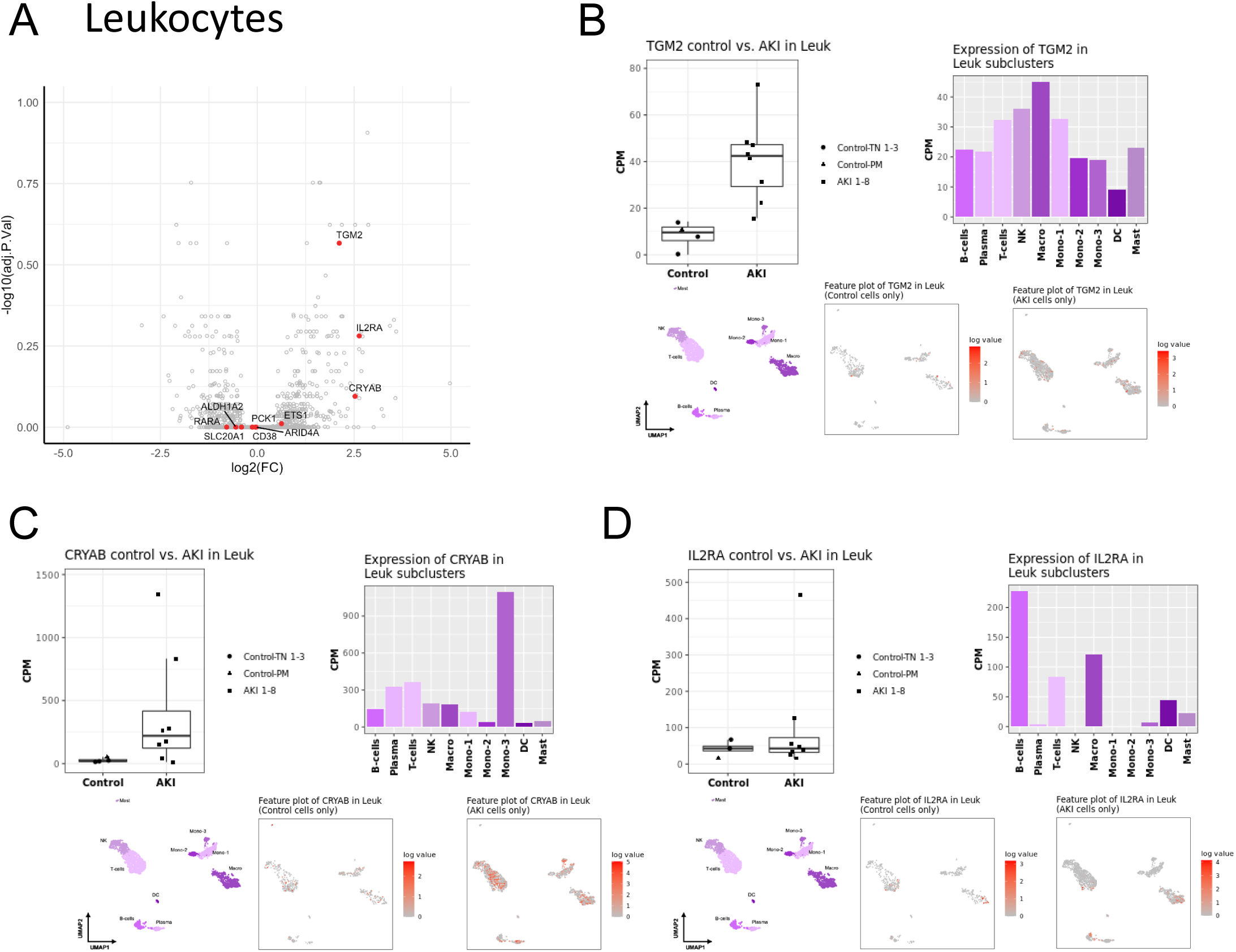
Expression of RAR target genes in renal leucocytes from patients with SA-AKI. **A,** Volcano plot indicating fold change in expression of core enrichment gene (AKI vs. controls) from Leukocyte snRNA seq dataset. Dotted line indicates p<0.05E. **B-D,** Expression of RAR core enrichment genes identified from GSEA analysis of the Leu snRNA seq dataset from Hinze et al. 2022. Figures were generated using on-line software https://shiny.mdc-berlin.de/humAKI/. Gene expression in controls vs. AKI patients in reads per million. UPMAP plots indicate leukocyte subsets identified from snRNA seq studies. Feature plots showing distribution of genes in leukocyte subsets summarized in bar charts.

**S. Figure 4.**
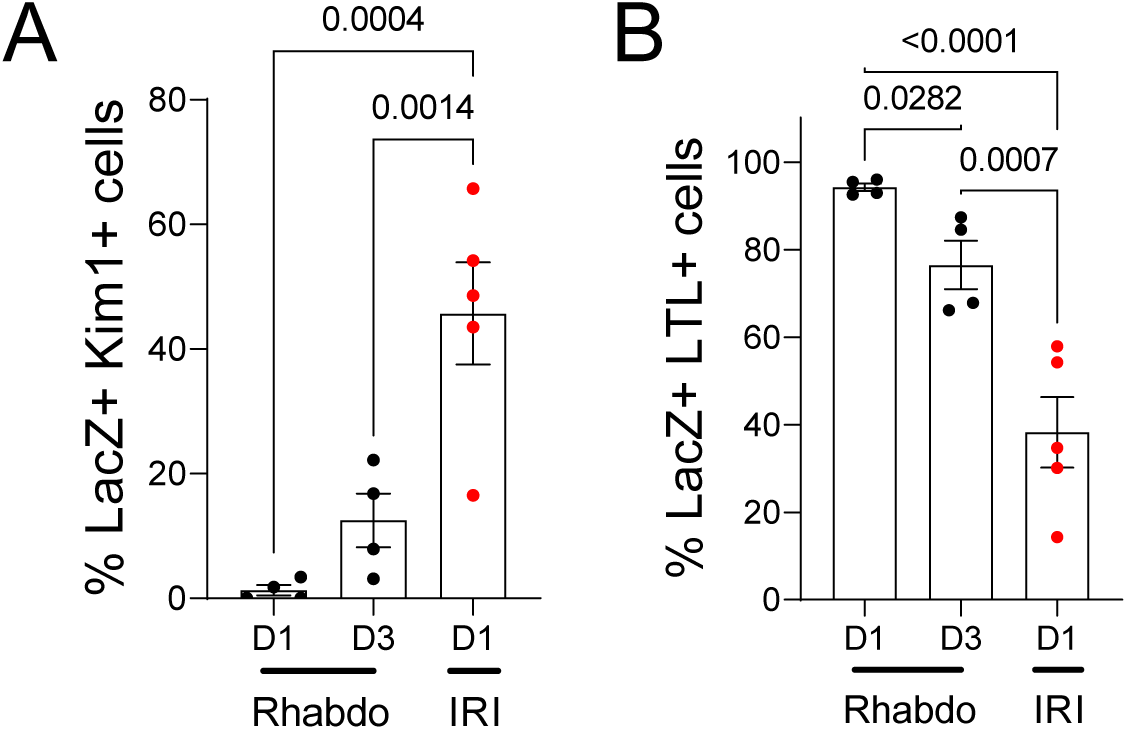
Increased activation of RAR signaling in injured, Kim1 positive PTECs after IRI compared with rhabdo-AKI. RARE-LacZ reporter mouse kidneys were used to evaluate the cellular localization of LacZ staining after IRI- and Rhabdo-AKI. IRI-AKI imaging data obtained from Chiba et al. 2016. ^2^ After LacZ staining, kidney sections were stained with LTL and antibodies to Kim1. LTL, Kim1 and LacZ staining evaluated in the OSOM. **A,** The % LacZ positive Kim1^+^ cells (injured PTECs); **B,** The % LacZ positive LTL^+^ Kim1^-^ cells (uninjured PTECs) at Day 1 and 3 after rhabdo-AKI, and Day 1 after IRI-AKI (maximal RARE LacZ activation after IRI-AKI^2^). Results expressed as mean +/- SEM, with individual data points shown. 1 way ANOVA used to compare between groups, q values shown for between group comparisons after correcting for repeated testing.

**S. Figure 5.**
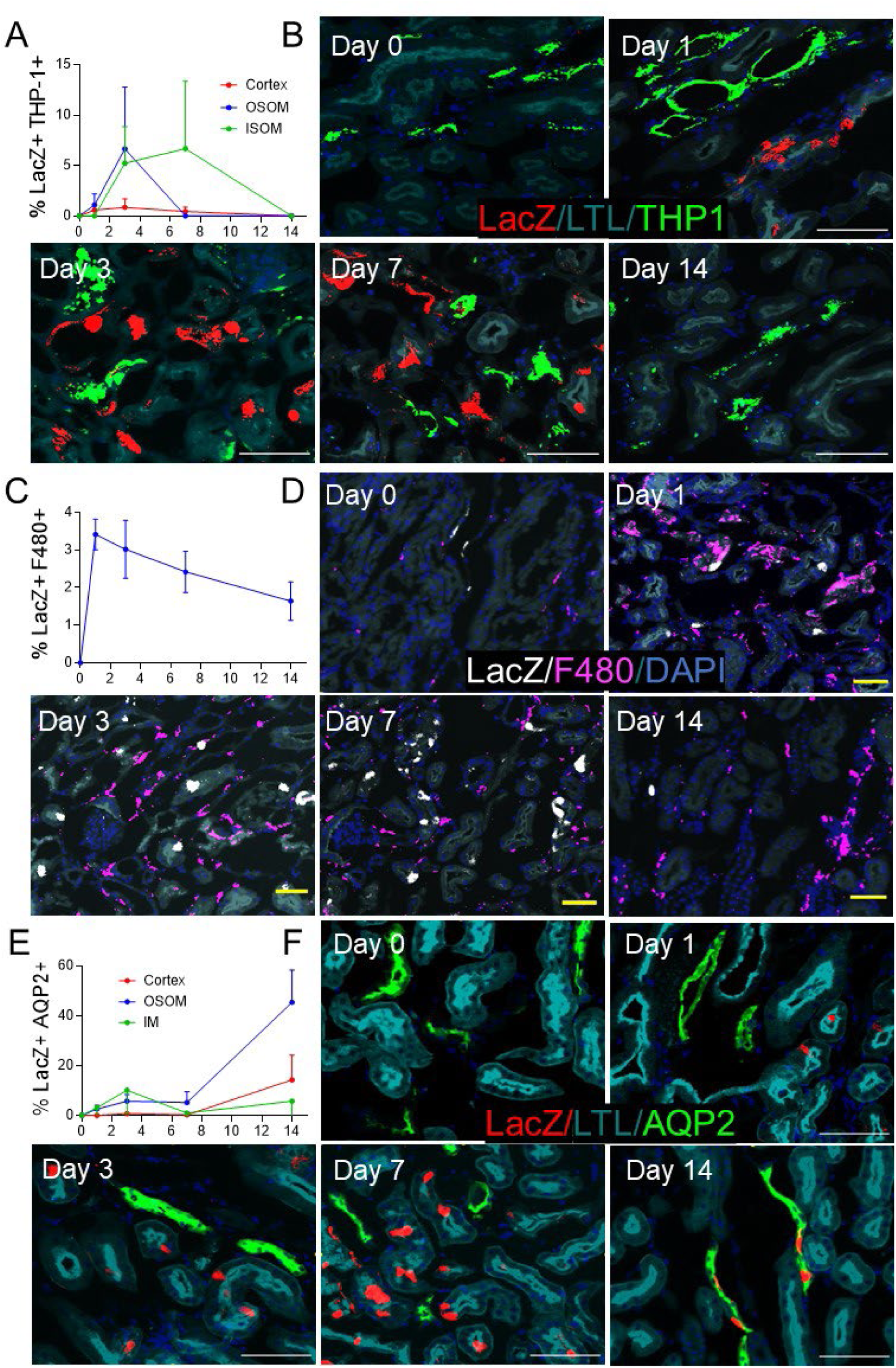
RAR signaling in thick ascending limb, renal macrophages, and collecting ducts after rhabdo-AKI. RARE-LacZ reporter mice were used to evaluate the cellular localization of RAR signaling after rhabdo-AKI. After LacZ staining, kidney sections were stained with LTL, Tam Horsfall Protein -1 (THP-1, thick ascending limb (TAL)), F4/80 antibodies (renal macrophages), and AQP2 (collecting duct (CD), principal cells). ***A/B, Activation of RAR signaling in TAL.*** The % LacZ positive THP-1^+^ TAL cells at different time points in the cortex, OSOM, and ISOM (A). Representative images showing LacZ, THP-1 and LTL staining in the OSOM (B). ***C/D, Activation of RAR signaling in renal macrophages.*** The % LacZ positive F4/80^+^ renal macrophages in the cortex and OSOM (C). Images showing LacZ, F4/80, and LTL staining in the OSOM (D). ***E, Activation of RAR signaling in CDs.*** The % LacZ positive AQP2^+^ CD cells at different time points after injury in the cortex, OSOM and IM (E). Images from the OSOM showing LacZ, AQP2, and LTL staining in the OSOM (F). Results expressed as means +/- SEM from 2 mice before injury (Day 0), and 4 mice each at Days 1, 3, 7 and 14 after injury. Scale bars B/F: 100μM; D: 50μM.

**S. Figure 6.**
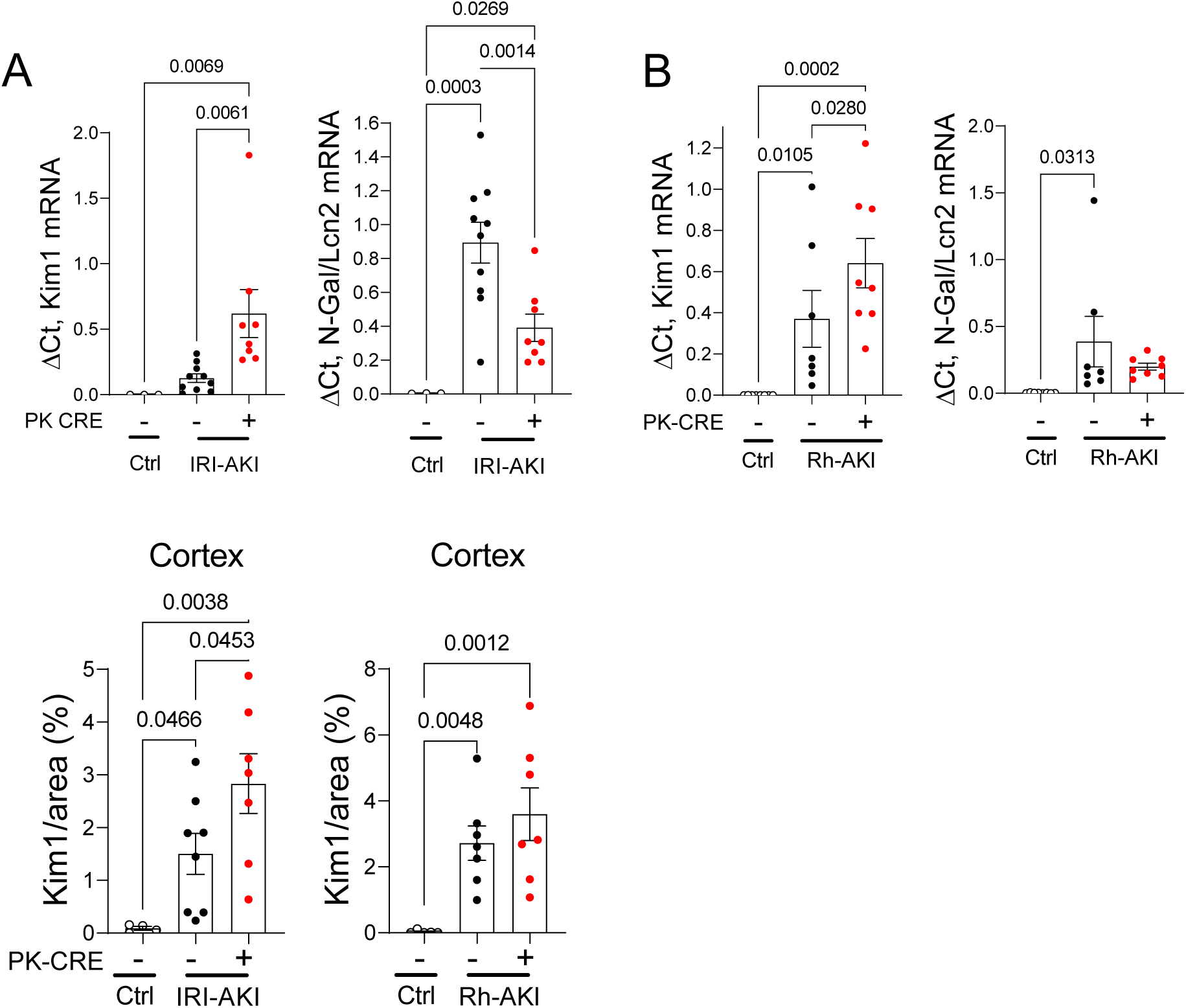
Expression of injury markers Kim-1 and N-Gal in the PTEC DN-RAR kidneys after IRI- and Rhabdomyolysis-AKI. PTEC DN-RAR mice underwent bilateral IRI-AKI, or rhabdo-AKI (Rh-AKI), and kidney harvested after 3 days. **A/B,** Renal *Kim-1* and *N-Gal* mRNAs by quantitative RT-PCR (QRT-PCR) after bilateral IRI-AKI, and rhabdomyolysis-AKI. **C,** Quantification of Kim-1 immunostaining as the % surface area in the cortex after IRI- and Rhabdo-AKI. Results expressed as means+/-SEM, with individual datapoints shown. 1-way ANOVA was used to compare between groups, and if p values <0.05, q values shown for between group comparisons after correcting for repeated testing.

**S. Figure 7.**
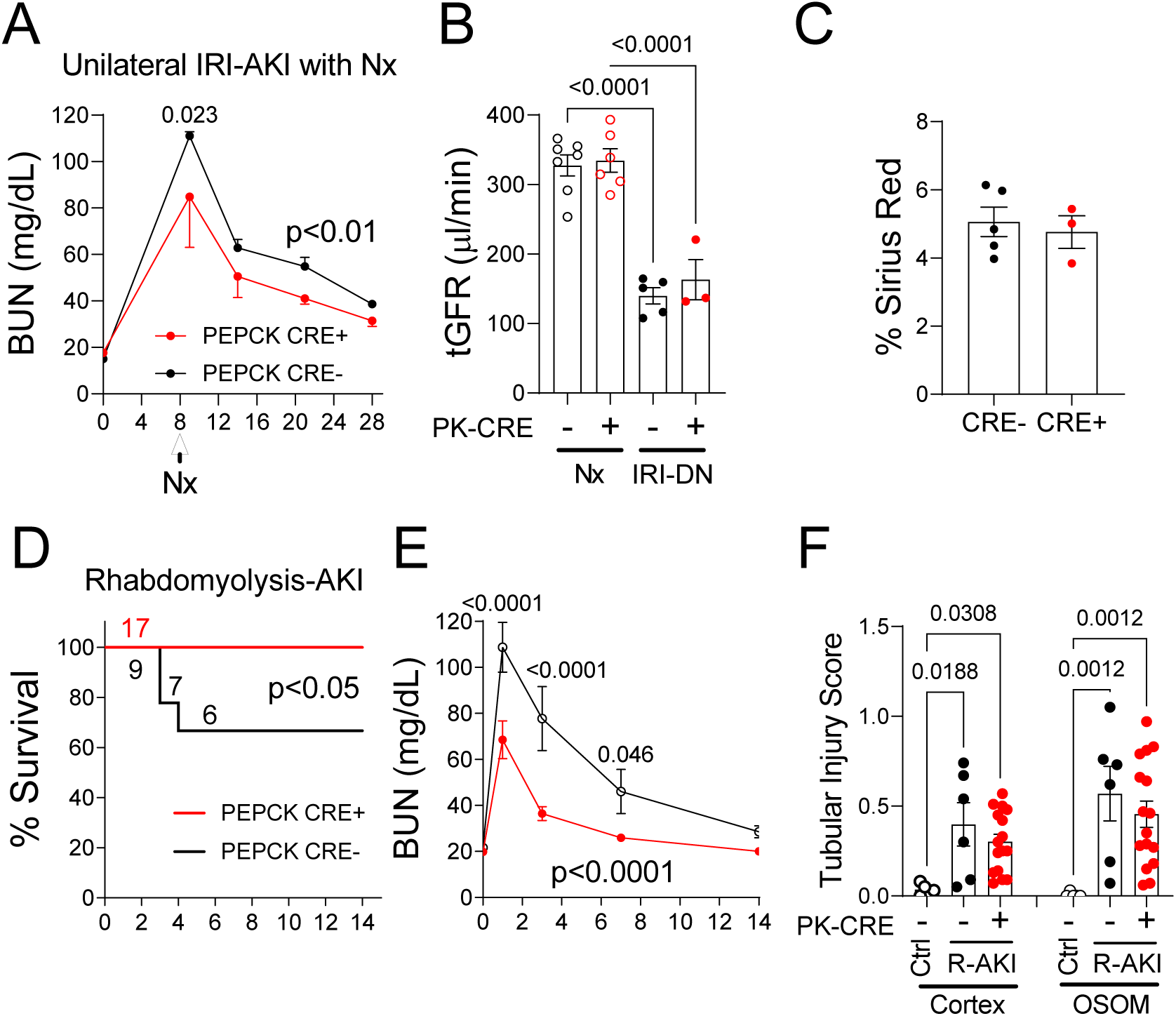
Long-term studies show that inhibition of RAR signaling in PTECs protects against AKI at early time points after IRI- and rhabdo-AKI but does not improve long-term functional or histological recovery. PTEC DN-RAR mice underwent unilateral renal pedicle clamping and delayed contralateral nephrectomy (Nx, unilateral IRI-AKI DN) and were followed up for 28 days after the initial injury, or rhabdomyolysis-AKI and were followed up for 14 days after injury. ***A-C, Unilateral IRI-AKI DN*. A,** BUN time course starting 1 days after Nx. Results expressed as means +/- SEM in mg/dl from 3 *PEPCK Cre+* and 5 *Cre-* controls. No mice died during these studies. **B,** Transdermal glomerular filtration rate (tGFR) corrected for body weight and expressed in μL/minute 26 days after injury. Results after injury (IRI-DN) compared with uninjured *Cre+* and *Cre-* controls 18 days after nephrectomy (Nx). Data points with mean +/- SEM are shown. C, Sirius red staining. Quantification as % surface area with Sirius red staining in the OSOM. ***D-F, Rhabdomyolysis-AKI.* D,** Survival curves. Mouse numbers indicated. E, BUN time course after injury, mouse numbers indicated in (D). F, Tubular injury scores. 14 days after injury in the cortex and OSOM in uninjured controls (Ctrl) and after rhabdo-AKI (R-AKI). A/E, 2-way ANOVA used to compare between groups over time. If p values <0.05 between groups, q values shown after correcting for repeated testing. B/C/F, 1-way ANOVA used to compare between groups, and if p values <0.05, q values shown for between group comparisons after correcting for repeated testing. D: Survival curves compared by Log-rank test, p<0.05.

**S. Figure 8.**
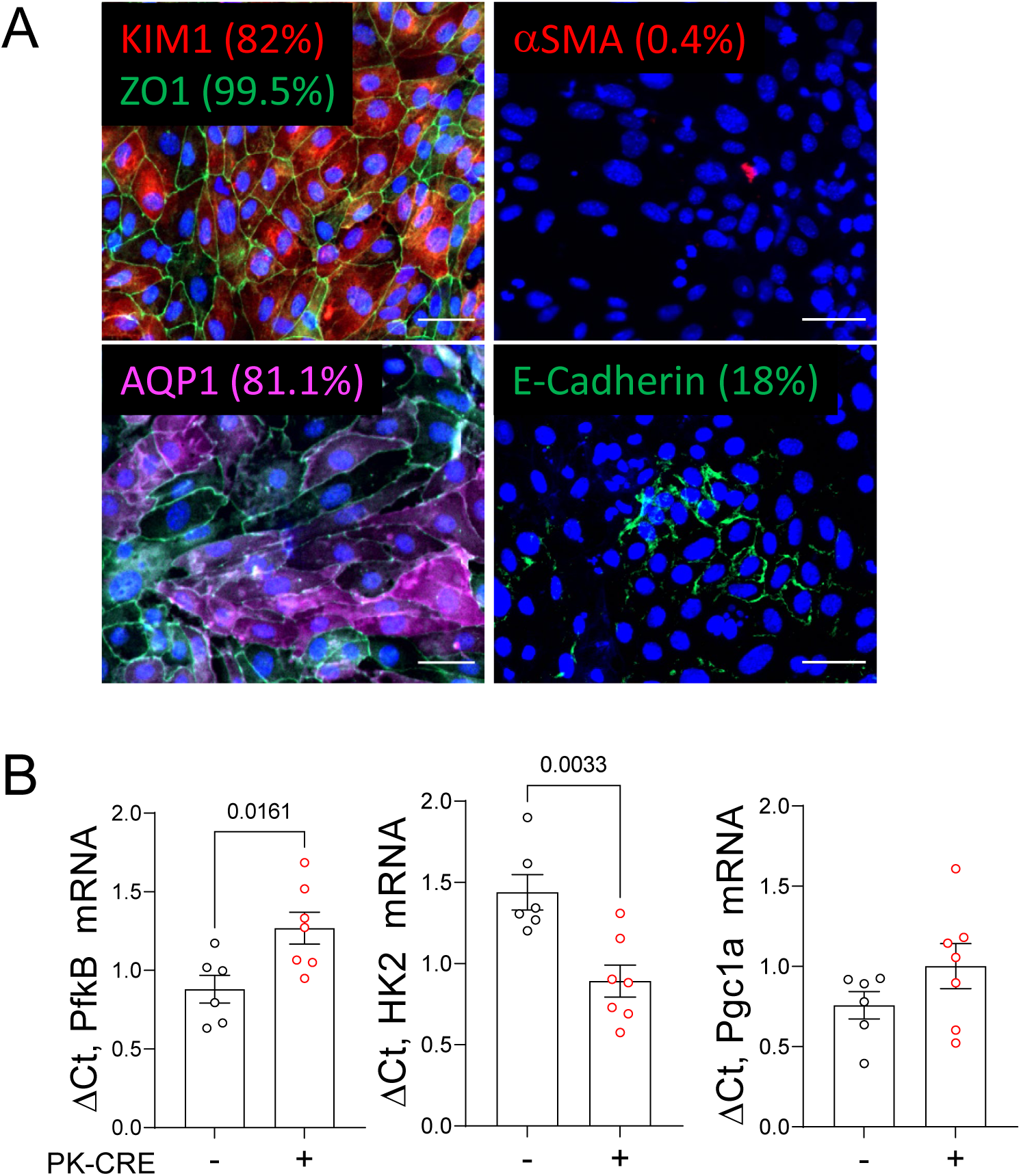
Characterization of primary PTECs isolated from PTEC DN RAR mice. Primary PTECs were isolated from homogenized and digested mouse kidneys and enriched using LTL-conjugated magnetic beads, as described in methods. **A,** Representative immunostaining of P2 PTECs Kim-1/ZO-1, AQP1, α- SMA, and E-Cadherin, with percentage of cells staining with the respective markers, as indicated. Scale bars=20μM. **B,** Quantification of *PfkB, Hk2,* and *Pgc1α* mRNAs by QRT-PCR in PTECs isolated from *PEPCK Cre+* and *Cre-* mice. Each data point is derived from RNA isolated from a separate P2 PTEC preparation from an individual mouse *Cre+* or *Cre-* mouse. Results are expressed as means +/- SEM, datapoints shown. T-tests were used to compare results in *PEPCK Cre+* and *Cre-* PTECs, p values shown if <0.05.

**S. Figure 9.**
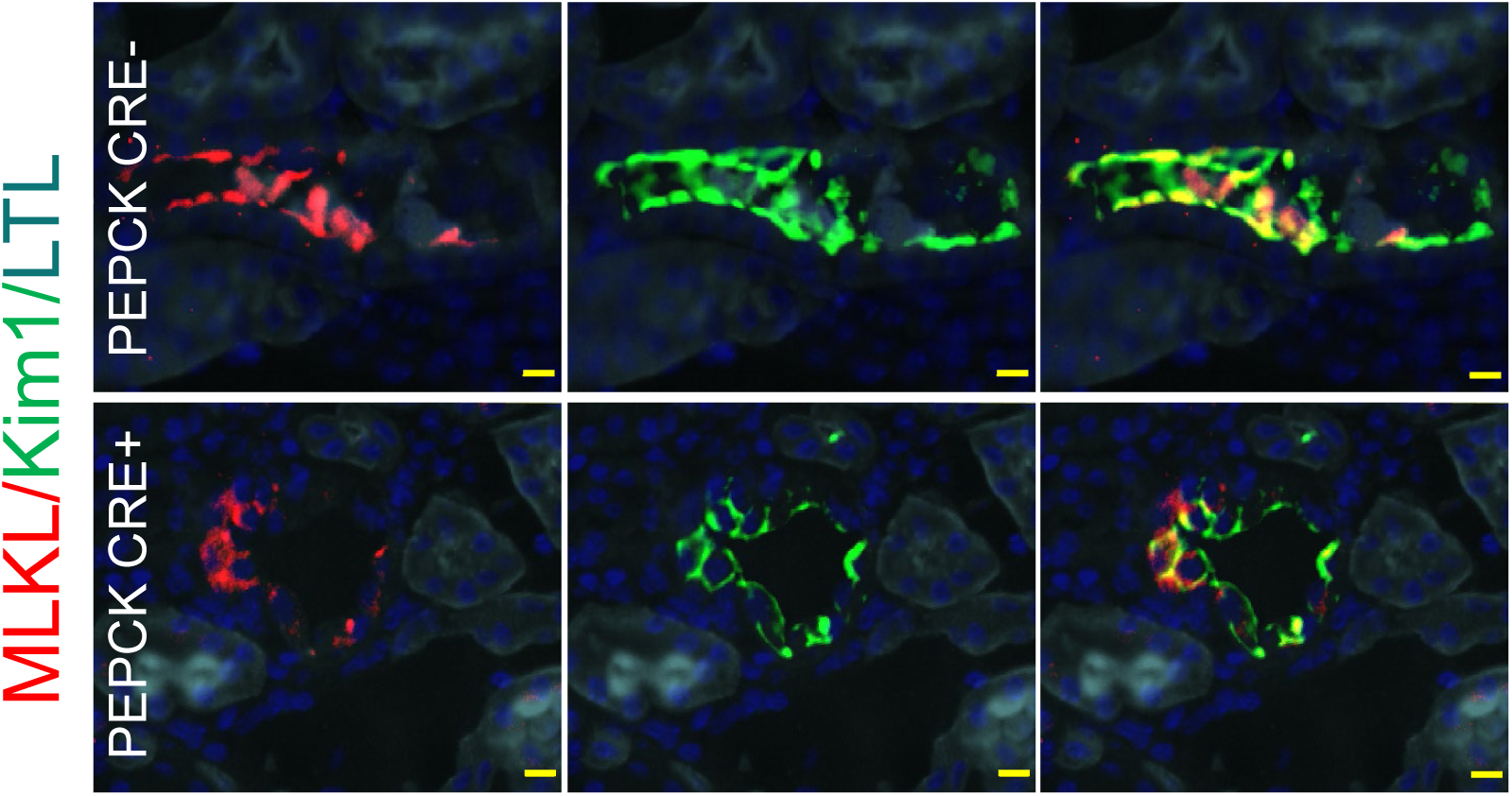
Subcellular localization of MLKL in *PEPCK Cre+* and *Cre-* mice after rhabdomyolysis AKI. PTEC DN-RAR mice underwent rhabdo-AKI, and kidneys were harvested after 3 days. Representative images showing MLKL and Kim-1 staining with dominant membrane localization of MLKL in Kim-1^+^ PTECs in *PEPCK Cre-* mice, and more diffuse cytoplasmic staining in Kim-1^+^ PTECs in *PEPCK Cre+* mouse kidneys after Rhabdo-AKI. Scale bars=10μM

**S. Figure 10.**
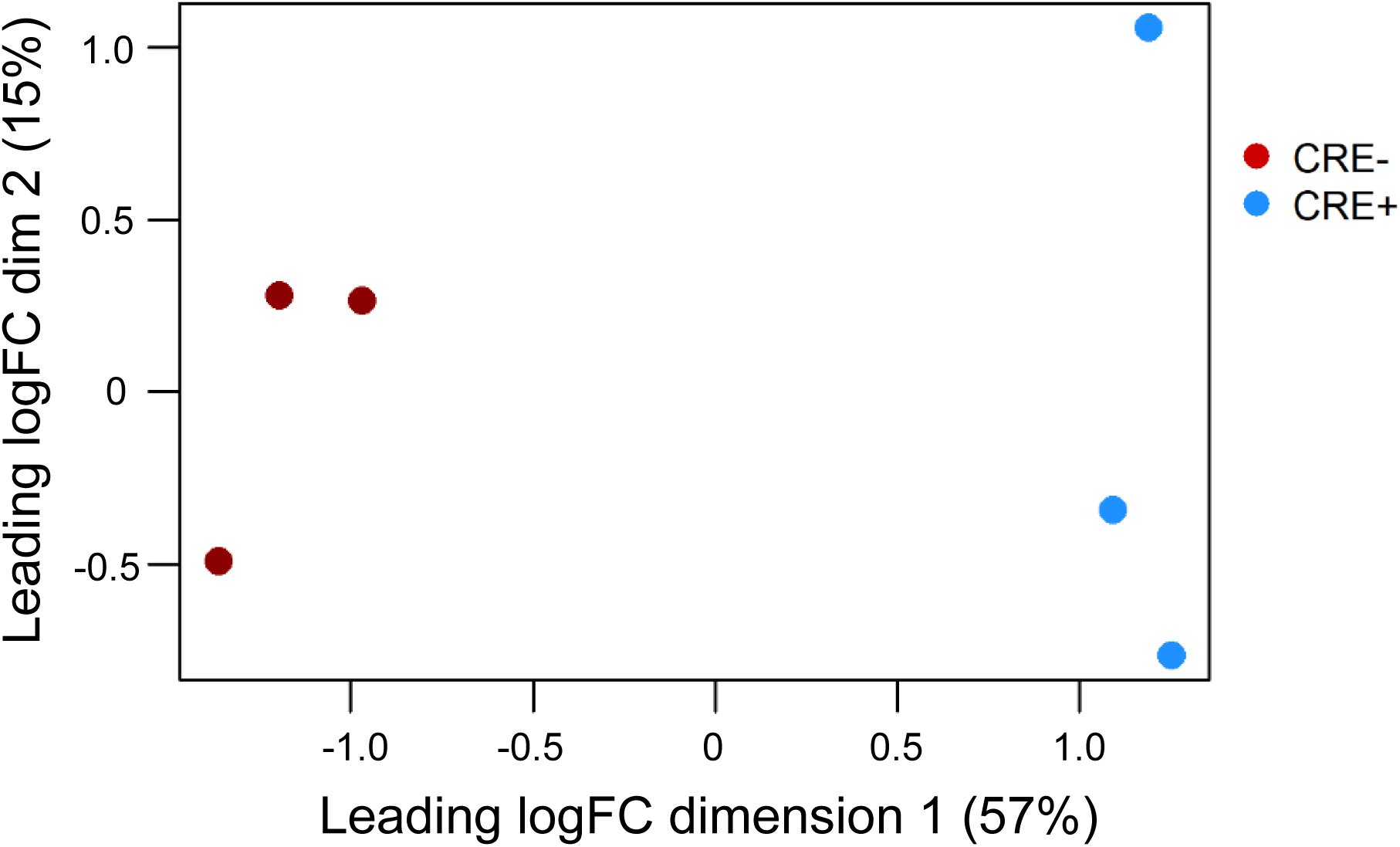
Principal component analysis (PCA) of CD11B+ RNA seq data. PTEC DN-RAR mice underwent bilateral IRI-AKI, and bulk RNA seq was performed on CD11B^+^ cells from 3 *PEPCK Cre+* and 3 *Cre-* mouse kidneys 3 days after injury. PCA shows clear separation of *Cre+* and *Cre-* datasets in the first dimension, which accounts for 57% of the variation.

**S. Figure 11.**
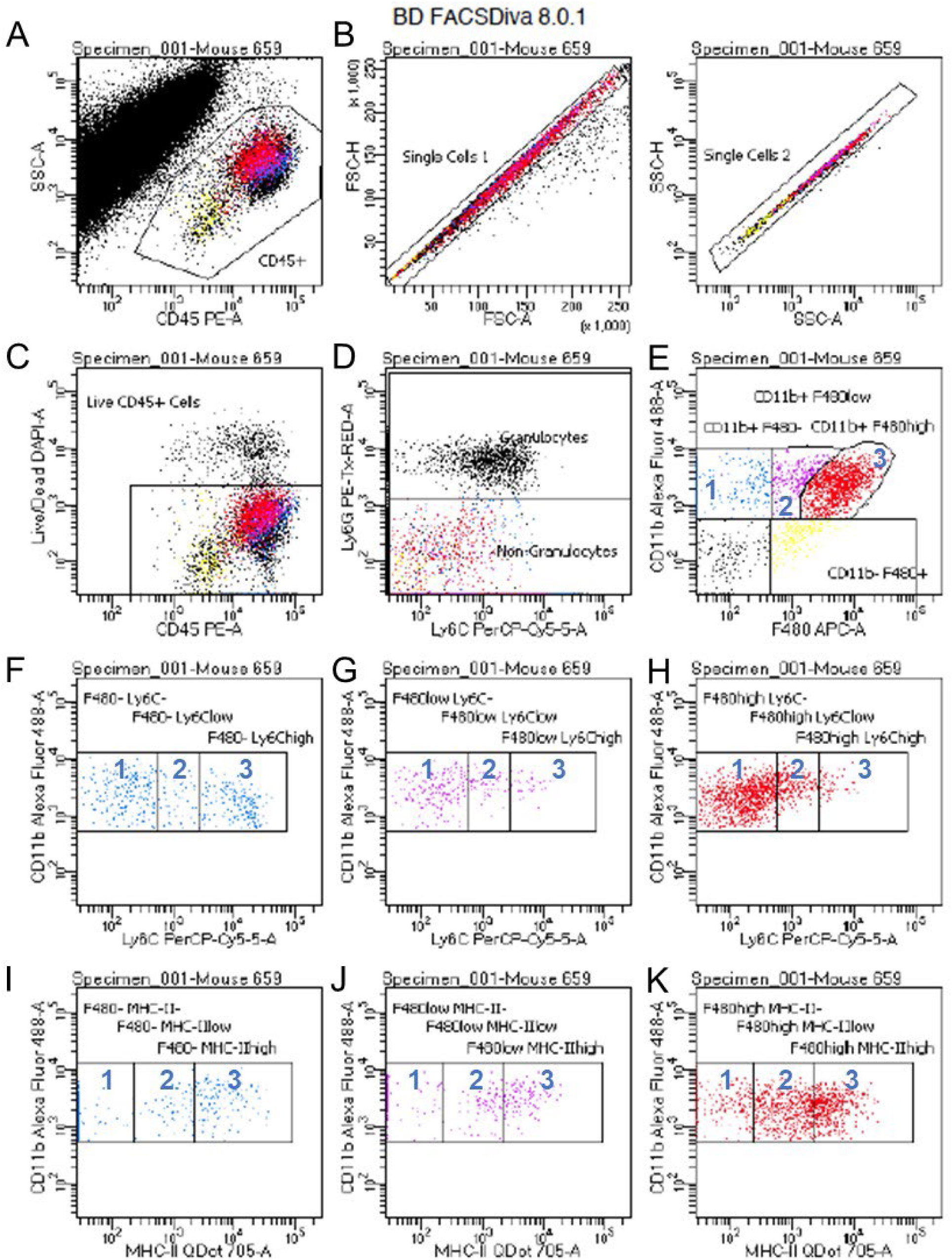
Flow cytometry gating strategy used to evaluate renal monocyte/macrophages. **A,** CD45^+^ cells: CD45 vs SSC-A; **B,** CD45^+^ single cells: FSC-H vs FSC-A, selected upper line followed by SSC-H vs. SSC-A select upper line; **C,** Single, CD45^+^ live cells: DAPI^-^ vs. CD45^+^; **D,** Exclude CD45^+^ granulocytes: Ly6G^+^ (granulocytes) and Ly6G^-^ cells (non-granulocytes); **E, *F4/80 expression in CD11B^+^; Ly6G^-^cells:*** F4/80^-^ (1); Intermediate (int), 2); high (hi) (3); **F-H, *Ly6C expression:* F,** F4/80^-^: Ly6C^-^ (1), Int (2), high (3); **G,** F4/80^Int^: Ly6C^-^ (1), Int (2), high (3); **H,** F4/80^Hi^: Ly6C^-^ (1), Int (2), high (3); **I-K, *MHC-II expression:* I,** F4/80^-^: MHC^--^ (1), Int (2), high (3); **J,** F4/80^Int^: MHC-II^-^ (1), Int (2), high (3); **K,** F4/80^Hi^: MHC-II^-^ (1), Int (2), high (3).

**S. Figure 12.**
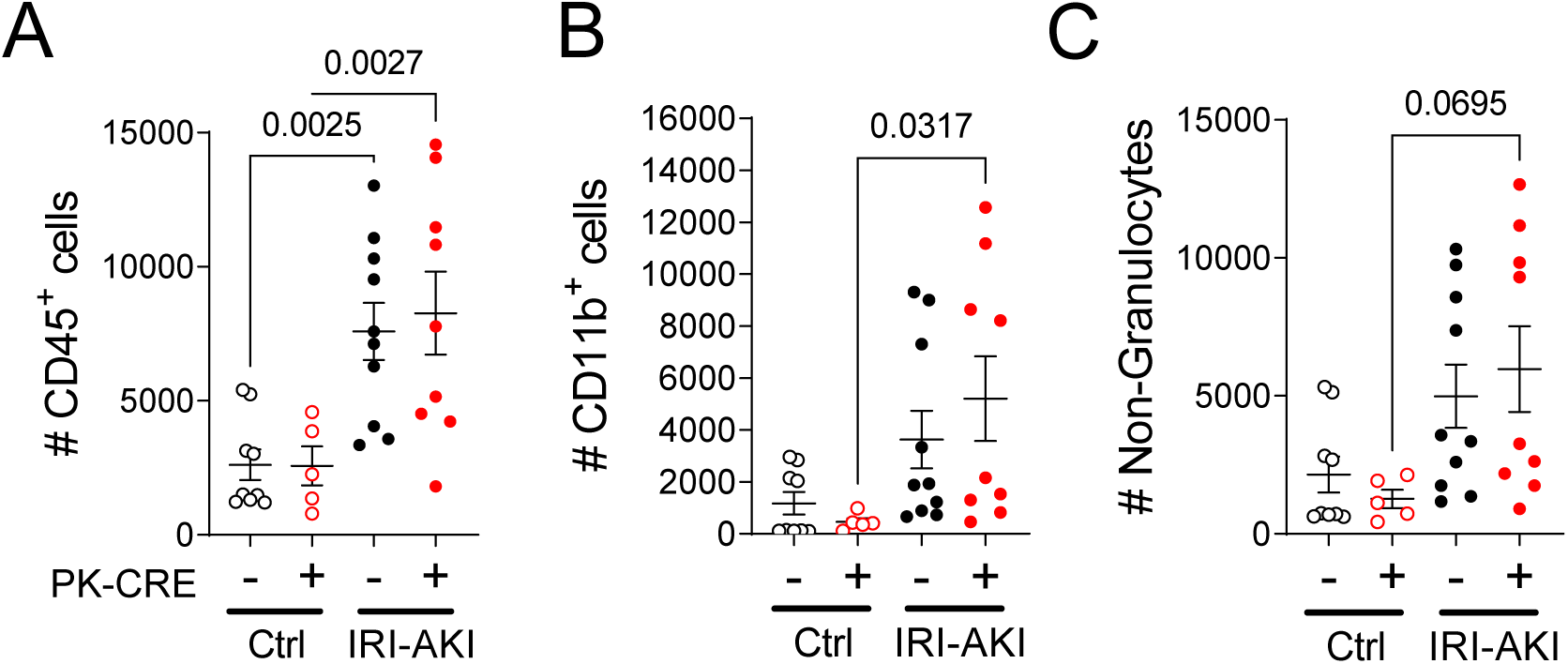
Absolute numbers of renal CD45^+^ and CD11B^+^ cells in PTEC DN RAR mouse kidneys. Kidneys were harvested from 5 *PEPCK Cre+* and 10 *Cre-* uninjured mice, and from 9 *PEPCK Cre+* and 10 *Cre-* mice 3 days after bilateral IRI-AKI. Tissue was digested and homogenized and evaluated by flow cytometry using antibodies and gating strategy illustrated in S. Fig. 11. **A,** Total numbers of live, single CD45+ cells acquired; **B,** CD11B+ cells; and **C,** non-granulocytes (CD11B^+^; Ly6G^-^). Results expressed as means +/- SEM, with individual data points shown. 1-way ANOVA was used to compare between groups, and if p <0.05, q values shown for between group comparisons, corrected for repeated testing.

**S. Figure 13.**
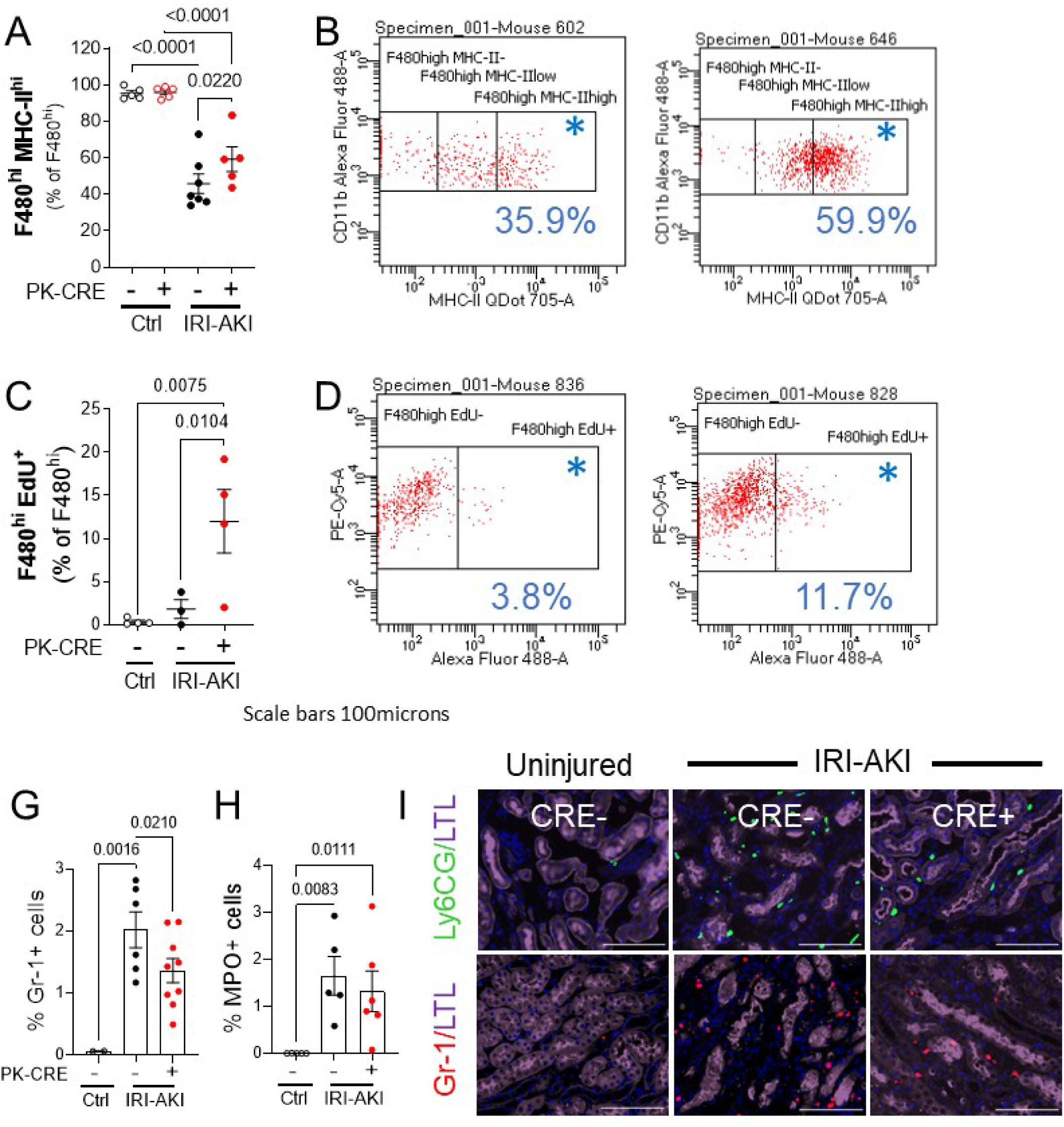
Flow cytometric and immunofluorescence analysis of renal monocyte/macrophages in PTEC DN RAR mice. Kidneys were harvested from PTEC DN RAR mice, uninjured and 3 days after bilateral IRI AKI. Tissue was digested, homogenized, and evaluated by flow cytometry. ***A/B, MHC-II^hi^ F4/80^hi^ cells (kidney resident macrophages).* A,** MHC-II^hi^ as the % of gated F4/80^hi^ cells. **B,** Representative CD11B and MHC-II expression chart in *PEPCK Cre+* and *Cre-* mice after IRI-AKI (% in gated area indicated with *). ***C/D, EdU^+^ F4/80^hi^ cells (proliferating kidney resident macrophages).* C,** EdU labeled cells as the % of gated F4/80^hi^ cells. D, Representative CD11B (Y axis) and EdU (X-axis) staining charts. ***E/F, Immunofluorescence staining with monocyte and neutrophil markers, Gr-1 and MPO.*** Kidneys were harvested from PTEC DN RAR mice 3 days after bilateral IRI-AKI and co-labeled with LTL and either Gr-1 (a marker of activated monocytes and neutrophils), or MPO (a marker of neutrophils) antibodies. **G/H, G,** Quantification of the Gr-1 and MPO+ cells as the % of the total cells counted. Results are expressed as means +/- SEM, with individual datapoints shown. 1-way ANOVA used to compare between groups, and if p <0.05, q values shown for between group comparisons, corrected for repeated testing.

